# Pangenome-aware DeepVariant

**DOI:** 10.1101/2025.06.05.657102

**Authors:** Mobin Asri, Pi-Chuan Chang, Juan Carlos Mier, Jouni Sirén, Parsa Eskandar, Alexey Kolesnikov, Daniel E. Cook, Lucas Brambrink, Glenn Hickey, Adam M. Novak, Lizzie Dorfman, Dale R. Webster, Andrew Carroll, Benedict Paten, Kishwar Shafin

## Abstract

Population-scale genomics information provides valuable prior knowledge for various genomic analyses, especially variant calling. A notable example of such application is the human pangenome reference released by the Human Pangenome Reference Consortium, which has been shown to improve read mapping and structural variant genotyping. In this work, we introduce pangenome-aware DeepVariant, a variant caller that uses a pangenome reference alongside sample-specific read alignments. It generates pileup images of both reads and pangenome haplotypes near potential variants and uses a Convolutional Neural Network to infer genotypes. This approach allows directly using a pangenome for distinguishing true variant signals from sequencing or alignment noise. We assessed its performance on various short-read sequencing platforms and read mappers. Across all settings, pangenome-aware DeepVariant outperformed the linear-reference-based DeepVariant, reducing errors by up to 25.5%. We also show that Element reads with pangenome-aware DeepVariant can achieve 23.6% more accurate variant calling performance compared to existing methods.

## Introduction

Human genome references are fundamental to genome analysis and research. Reference genomes provide a stable coordinate system to analyze and compare genomic variation among samples^1–3^. A single linear genome sequence as a reference genome like GRCh38 has been the primary mode of reference-based genomic analysis^4^. However, reference bias can occur in locations where the reference diverges sufficiently from a sample under study^5^. This hinders accurate mapping and variant calling, which can mislead downstream analyses^6–8^. One way to address reference bias is using a pangenome reference, which includes diverse, non-redundant DNA sequences from multiple individuals typically embedded in a graph structure^9,10,11^.

Recently, the Human Pangenome Reference Consortium (HPRC) assembled 88 haplotypes from 44 individuals and integrated them into a pangenome graph^12^. This graph was then used with pangenome-aware read mappers to map short reads to the GRCh38 reference. By providing these mappings to DeepVariant^13^, the number of calling errors reduced by 34% compared to linear-reference-based pipelines^12^. The proprietary software DRAGEN also claims to remove nearly half of the errors by using a pangenome although how the pangenome is used and the sequence contents of the pangenome have not been publicly disclosed^14^.

Population information can enhance variant calling not only by improving read mapping but also by being directly integrated into the calling process. The GATK HaplotypeCaller “--population-callset” option^15,16^ and an allele-frequency-based DeepVariant model^17^ use population allele priors to adjust genotype calls. However, as these approaches are frequency, not pangenome-based, they cannot use long-range haplotype information. Pangenome-based genotypers can call complex and structural variants with k-mer mapping approaches^18,19^ but the use of k-mers instead of more sophisticated models which model read-level errors limits their accuracy of small variants, variants not in the graph, and repetitive regions.

In this work, we present pangenome-aware DeepVariant, which uses a Convolutional Neural Network (CNN) to jointly model sample-specific reads and pangenome sequences, enabling more accurate variant calling than using read-level information alone. We train with multiple samples and short read sequencing platforms to generalize across ancestries and instruments. We show that using pangenome information improves accuracy in samples mapped to a linear reference with BWA-MEM^20,21^, and larger accuracy improvements in samples mapped by vg giraffe^8^. Although pangenome-aware DeepVariant uses the pangenome to improve accuracy, it produces final calls in the coordinates of a linear reference, such as GRCh38, enabling direct use in downstream methods based on GRCh38.

We also explored running pangenome-aware DeepVariant with personalized pangenomes, which subsample the pangenome to the sequences most relevant to the sample^22^. As the pangenome references increase in the number of haplotypes, the ability to subsample will become essential to preserve computational scalability. In this work, we show that personalized pangenomes can achieve nearly the same accuracy as the full pangenome.

## Results

### Pangenome-aware DeepVariant Overview

Pangenome-aware DeepVariant (DV) calls sample-specific short variants (<50 bp) using reads mapped to a linear reference (e.g., GRCh38), along with a pangenome reference to improve the inference accuracy. The process begins by identifying candidate variant sites from read alignments. For each candidate site, it creates a 221 bp image centered on the variant, showing various features of the read and pangenome haplotype alignments (**Methods, Supplementary Figure 1**). To generate these pileup images, haplotype alignments must be extracted from the pangenome; therefore, the pangenome must include the linear reference to which the reads were mapped. The images are then passed through a Convolutional Neural Network (CNN) that classifies each candidate to have a genotype either of reference allele, heterozygous variant or homozygous alternate variant. Finally the model’s output is post-processed to derive a VCF file containing the genotypes for the passed variants.

For training the CNN model we created a training dataset consisting of Illumina and Element reads from 6 different samples, mapped to the GRCh38 reference using both BWA-MEM^20,21^ and vg giraffe^8^. The true genotypes were labeled using three different truth sets; Genome-In-A-Bottle-v4.2.1 for HG004-HG007 samples^23^, T2T-Q100-v1.1 for HG002^24,25^, and the Platinum Pedigree for HG001^26^ (**Supplementary Table 1**). We evaluated pangenome-aware DV using the v1.1 pangenome graph released by the HPRC, which includes 88 reference-quality assembled haplotypes^12^ (**Figure 1**). We also generated personalized genomes using the vg toolkit^22^, which can be alternatively used as the input pangenome (**Methods, Figure 1**).

**Figure 1:**
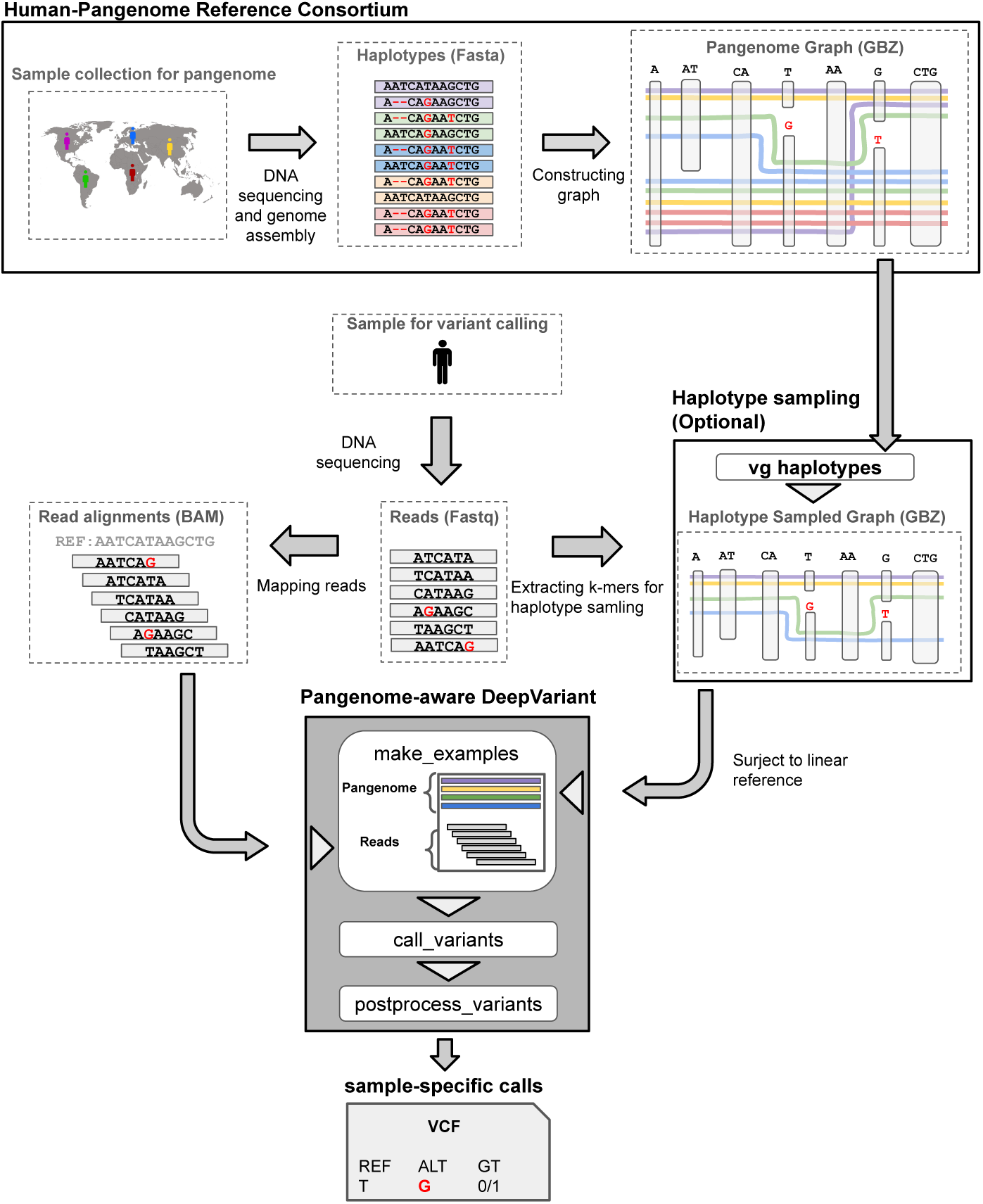
Overview of pangenome-aware DeepVariant. The HPRC created a pangenome graph using a genetically diverse set of the assembled haplotypes. This graph or a personalized version is fed to pangenome-aware DeepVariant to augment its read pileup images with the related pangenome haplotypes. It then uses the pangenome-augmented pileup images to infer sample-specific variant calls. In the example provided in this figure, the true variant is heterozygous (T/G) with a weak signal in short reads, leading to low confidence in variant calling without using a pangenome. However, due to the predominance of the alternative allele “G” in the human population, pangenome-aware DeepVariant can call this variant with higher confidence.

### Benchmarking Illumina and Element calls against T2T-Q100 and Platinum truth sets

We performed variant calling with Illumina NovaSeq reads (with 35x coverage) for the HG002 sample and benchmarked the output calls against the T2T-Q100-v1.1 truth set. This truth set was derived from the high-quality diploid assembly released by the T2T consortium, providing a more complete and accurate call set compared to the conventional truth sets like GIAB-v4.2.1 (more details in **Methods: Truth sets and training dataset**). Using linear-reference-based DV with the BWA-MEM read mappings, we achieved 99.32% precision and 97.91% recall, with 125,309 total errors. Using pangenome-aware DV with the full HPRC pangenome (88 haplotypes) and the same read mappings reduced the number of errors to 119,050, a 5% improvement over linear-reference-based DV (**Figure 2a**). Using vg giraffe mappings we obtained almost the same amount of error reduction (5.1%) when we switched from linear-reference-based DV to pangenome-aware DV. It shows that regardless of how the reads are mapped, providing a pangenome is useful for inferring correct variants. Since the T2T high-confidence bed file contained the regions not necessarily mappable by short reads we created a more conservative bed file by intersecting T2T-v1.1 and GIAB-v4.2.1 high-confidence bed files, which covered about 80% of the genome. Limiting our analysis to this bed file we observed 11.2% and 13.5% reductions in errors when using pangenome-aware DV with BWA-MEM and vg giraffe mappings respectively (**Supplementary Table 18**). The improvements made by pangenome-aware DV were more than 95% driven by SNPs, using either vg giraffe or BWA-MEM mappings (**Supplementary Table 4**).

**Figure 2:**
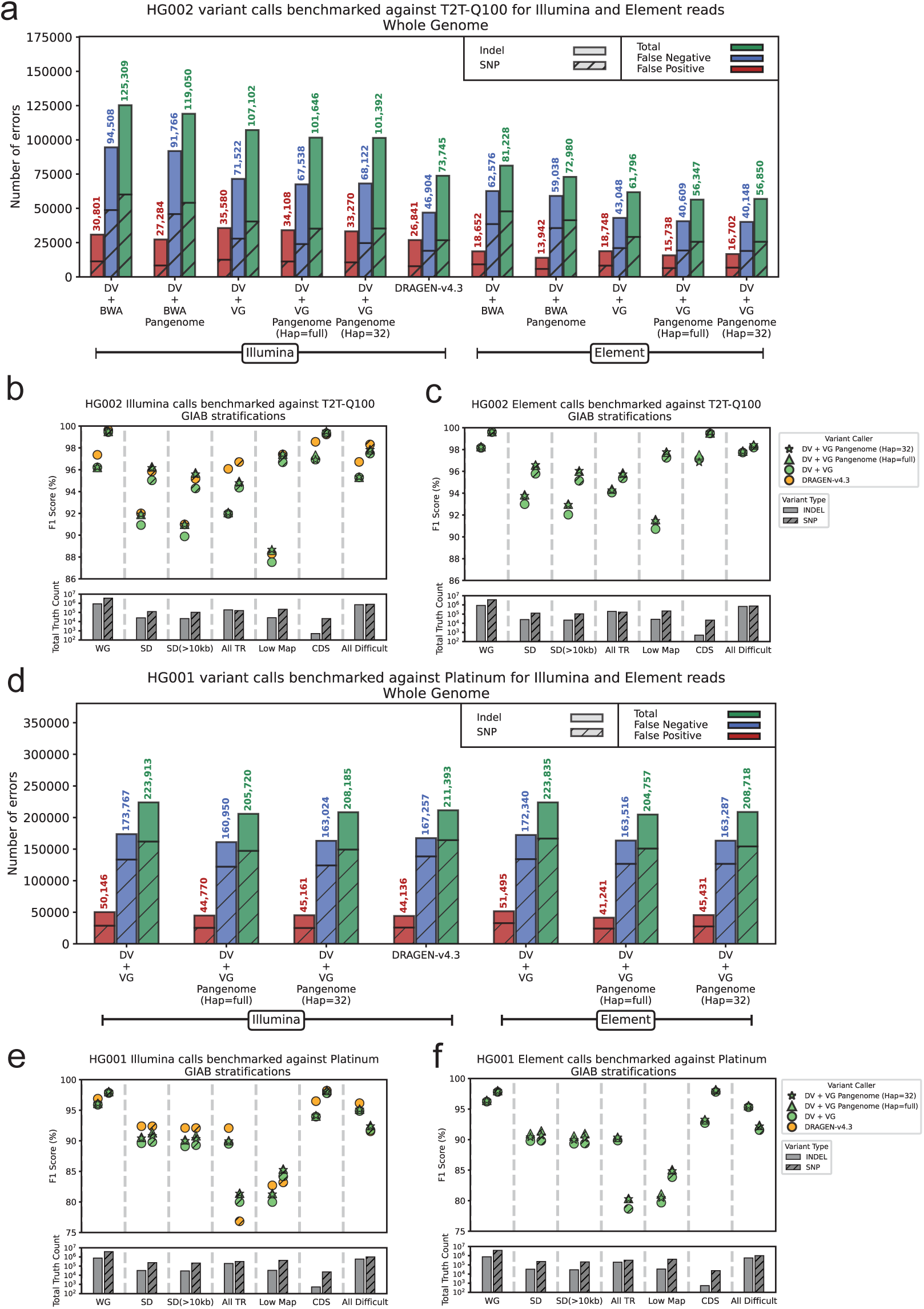
Benchmarking Illumina and Element calls against T2T-Q100 and Platinum truth sets . **a)** HG002 Illumina and Element calls are benchmarked against the T2T-Q100 truth set across the T2T high- confidence regions. Different modes of DeepVariant (DV) have been tested with vg giraffe and BWA- MEM. The x-axis labels with “Pangenome” refer to the pangenome-aware DeepVariant and for the rest, the linear-reference-based DeepVariant was used. The x-axis labels with “(Hap=full)” refers to using all 88 haplotypes in the HPRC-v1.1 pangenome and “(Hap=32)” refers to using a personalized pangenome with 32 haplotypes. **b)** Performance of variant calling across seven GIAB stratifications. The x-axis labels are “WG” (whole genome), “SD” (segmental duplications), “SD (>10kb)” (segmental duplications longer than 10 kb), “All TR” (all tandem repeats), “Low Map” (regions with low mappability), “CDS” (protein coding DNA sequence), “All difficult” (all regions difficult for variant calling). At the bottom of the panel the total number of SNP and indel truth variants existing for each stratification is shown. **c)** Similar to (b) but for Element calls. **d, e and f)** Similar to (a), (b), and (c) respectively but for the HG001 calls benchmarked against the Platinum truth set. (the whole-genome high-confidence bed file is also changed to the one specific for Platinum)

Our experiments with Element Biosciences data (with 40x coverage) additionally supported the benefit of pangenome-aware DV for variant calling. For example, running pangenome-aware DV with the full pangenome and the Element reads mapped with vg giraffe, we observed 8.8% reduction in errors compared to linear-reference-based DV (from 61,796 to 56,347 total errors) (**Figure 2a**). For each specific combination of mapper and variant caller, the call set created with Element data was more accurate than the one created with Illumina. This is consistent with the results from other studies showing that Element data has improved base-level accuracy compared to Illumina, especially in homopolymer runs and short tandem repeats^27^. Among all our assessments against the T2T-Q100 truth set, the most accurate call set for short reads was the output of mapping Element reads with vg giraffe and running pangenome-aware DV with the full pangenome (achieving a precision of 99.65% and recall of 99.1%) (**Figure 2a, Supplementary Table 5**).

We also tested pangenome-aware DV with a personalized pangenome (32 haplotypes) and found its performance to be nearly identical to that with the full pangenome (88 haplotypes) (**Figure 2a and 2d**). This demonstrates that pangenome-aware DV is compatible with personalized pangenomes, a key advantage when the original pangenome is too large to embed entirely into the input pileup image (**Methods: Creating pileup images augmented with pangenome**).

We also performed variant calling with some external callers such as Octopus, GATK (with and without DRAGEN submodules), and DRAGEN-v4.3 (**Methods**). For GATK and DRAGEN we used their internal mapping pipelines. We ran Octopus with both BWA-MEM and vg mappings and observed 11.6% reduction in total errors when vg mappings were used, similar to our results with DeepVariant. DRAGEN-v4.3 had the best performance with 73,745 total errors, which is still 30% more errors compared to our best configuration for pangenome-aware DV with 56,347 errors (**Figure 2a**). Noticeably, DRAGEN is a proprietary software for Illumina which uses its own pangenome and mapping pipeline^14^. DRAGEN is able to optimize its accuracy specifically for Illumina platform whereas Pangenome-aware DeepVariant provides a more generalizable solution for multiple platforms, such as Element. The pangenome incorporated in DRAGEN is not publicly available, therefore quantifying DRAGEN against pangenome-aware DV with the exact same pangenome has not been possible.

Benchmarking against the Platinum truth set, the Illumina-based calls inferred by linear-reference-based DV had 223,913 total errors (98.88% precision and 96.21% recall rates) (**Figure 2d**). Using pangenome-aware DV with full and personalized pangenomes decreased the errors by 8.12% and 7.0% respectively, aligning with the T2T-Q100 benchmarking results. The Platinum truth set generation was heavily dependent on Illumina which could be the reason we don’t see an improvement with Element data as we observed with the T2T-Q100 truth set. Additionally we compared DRAGEN versus pangenome-aware DeepVariant using Platinum truth set. Compared to Illumina-based DRAGEN, which had 211,393 total errors, pangenome-aware DV—regardless of whether it used Illumina or Element data, or full or personalized pangenomes—had 1.2% to 3.1% fewer errors (**Figure 2d, Supplementary Tables 6 and 7**).

To identify which parts of the genome are most impacted by variant calling with pangenomes, we stratified T2T-Q100 benchmarking results using GIAB-v4.2.1 stratification bed files.

Segmental duplications and regions with low mappability were among the stratifications with the highest level of improvement. Using pangenome-aware DV with the full pangenome and Illumina data mapped with vg giraffe we achieved 97.36% precision and 93.76% recall in segmental duplications, with 20.3% and 6.92% fewer errors compared to linear-reference-based DV and DRAGEN-v4.3 respectively (**Supplementary Table 4, Figure 2b**). Similar to the whole-genome results the best performance in segmental duplications was obtained by mapping Element reads with vg giraffe and variant calling with pangenome-aware DV, resulting in 97.87% precision and 94.41% recall. (**Supplementary Table 5, Figure 2c**). Stratification analysis was performed similarly for the Platinum truth set and the improvements enabled by the use of pangenomes were still apparent in segmental duplications and regions with low mappability (**Supplementary Tables 6 and 7**, **Figure 2e and 2f**).

To ensure that our models are not overfitted to the chromosomes included in the training dataset, we benchmarked HG002 calls separately in chromosome 16, which was held out during training. Using different combinations of sequencing platforms and read mappers, we could achieve up to 6.4% reduction in errors compared to linear-reference-based DV in chromosome 16 (**Supplementary Figure 16, Supplementary Tables 8 and 9**). We also performed variant calling for the HG003 sample, which was excluded from the training dataset. We used both Illumina and Element reads for variant calling and benchmarked the output calls against the GIAB-v4.2.1 truth set since the T2T truth set was not available for this sample. The results were consistent with the previous T2T-based assessments showing up to 25.5% reduction in errors using pangenome-aware DV and supporting the advantage of our method even for the samples not used in model training (**Supplementary Figures 17 and 18, Supplementary tables 2 and 3**).

### Investigating SNP errors fixed or induced by pangenome-aware DV

In our observation, pangenome-aware DeepVariant (DV) significantly improved variant calling in segmental duplications (SDs). For more investigation we analyzed the SNP call improvements and classified them as either “removed FP” (false positives removed by pangenome-aware DV) or “rescued FN” (false negatives rescued by it), using HG002 Element data and the T2T-Q100 truth set. We focused this analysis only on SNPs which were the primary drivers of the observed improvements. We also partitioned the whole genome high-confidence bed file into 4 regions: SDs with identity greater than 99% (covering 12.23 Mb of the genome), SDs with identity between 99% and 98% (16.08 Mb), SDs with identity between 98% and 95% (23.98 Mb),and the rest of the genome (2687.37 Mb). SDs with identity less than 95% were added to the partition of “the rest of the genome”. In our genome-wide analysis of 9,272 improved SNP sites, we observed the total number of removed FP (4,462) is nearly equal to the total number of rescued FNs (4,810) showing that pangenome is beneficial for both types of accuracy improvements (**Figures 3d, Supplementary Table 13**).

**Figure 3:**
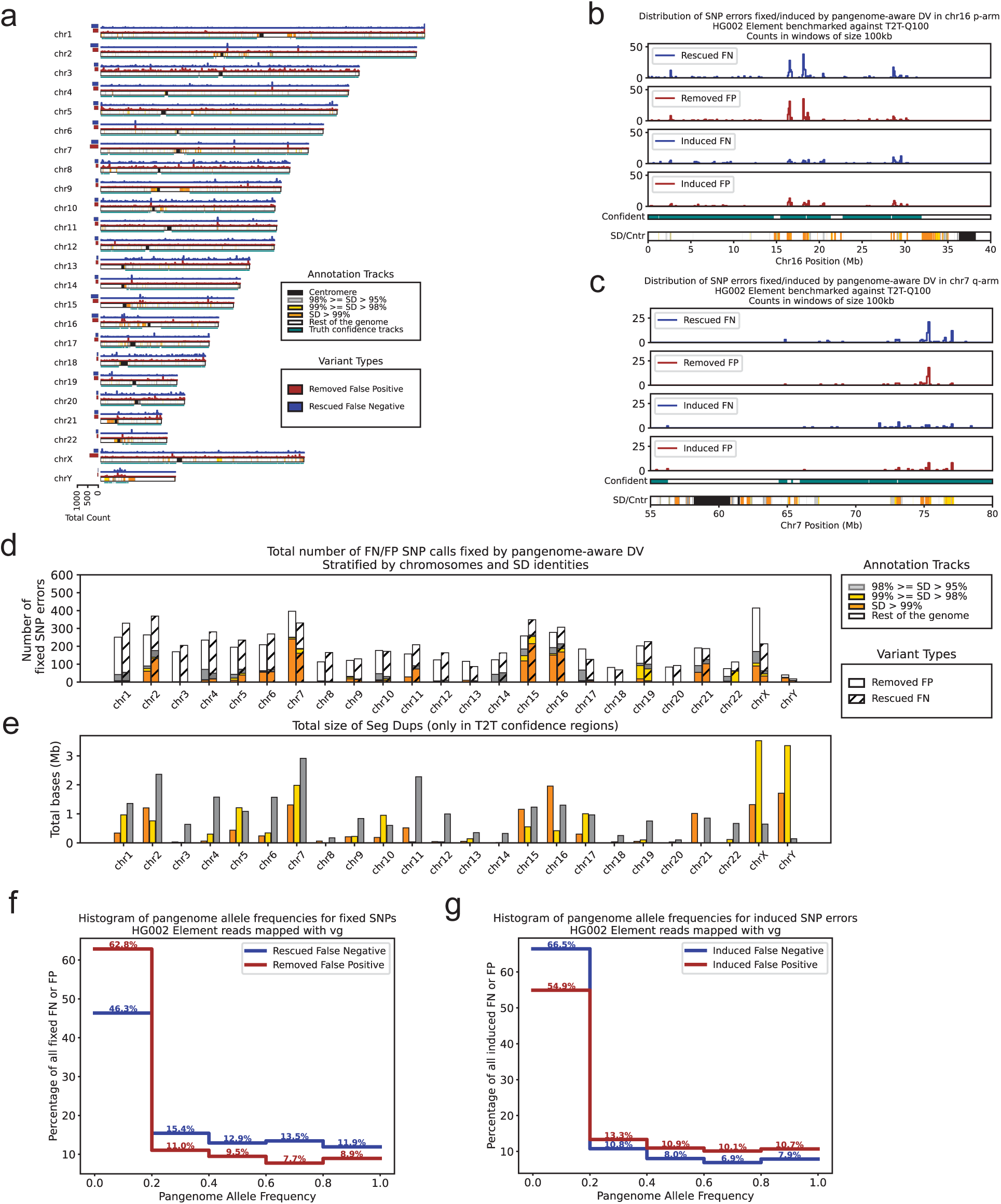
Investigating SNP errors fixed or induced by pangenome-aware DV. All the panels in this figure are based on variant calling with HG002 Element reads that were mapped with vg giraffe and the truth set for benchmarking was T2T-Q100-v1.1. **a)** The counts of the fixed SNP errors in 100kb windows are plotted across the GRCh38 chromosomes. The y axis for each chromosome is scaled separately.The removed FP and rescued FN calls are shown with red and blue respectively. The bottom track for each chromosome is showing 5 different annotations; centromere (black), SDs with identity greater than 99% (orange), SDs with identity between 99% and 98% (yellow), SDs with identity lower than 95% (light gray) and the rest of the genome (white). Below the SD/Cntr annotation the tracks with teal color show the high-confidence regions for the T2T-Q100 truth set. The left horizontal barplots show the total counts of fixed variants per chromosome. **b)** Distribution of fixed and induced errors are shown in the p-arm of chr16. The number of variants are counted in adjacent non-overlapping windows of length 100kb. The bottom annotation and high-confidence tracks are colored similar to panel (a). **c)** Similar to panel (b) but for the q-arm of chr7 **d)** The total number of variants fixed by pangenome-aware DV across different chromosomes stratified by SD identities and centromeric regions. If there was an overlap between SD and centromere they were labeled as centromere. The bars are colored similar to panel (a). The bars for removed FP (solid) and rescued FN (hashed) are plotted separately. **e)** The total size of segmental duplications across different chromosomes, which are colored by identity similar to panel (a). **f)** The frequency of HPRC alleles are binned with the size of 0.2 on the x axis. Each fixed variant is associated with a HPRC allele frequency extracted from the raw HPRC vcf file and the number of fixed variants are counted in each frequency bin. **g)** Similar panel (f) but for errors induced by pangenome-aware DV.

When we measure the impact of improvements by looking at the rate of fixed SNP per Mb, we observe that “SDs with identity greater than 99%” had the highest rate of fixing errors with 74.82 SNP/Mb for removed FP and 86.26 SNP/Mb for rescued FN (161.08 SNP/Mb fixed in total) (**Figures 3d and 3e, Supplementary Table 15**). This shows the effectiveness of pangenome-aware DV in highly duplicated regions. For example, **Figures 3b and 3c** illustrate the q-arm of chromosome 7 and the p-arm of chromosome 16, showing how induced and fixed errors are distributed across their segmental duplications. They show that almost all the segmental duplications present in the high-confidence regions are enriched with fixed errors. However the current results might not reflect the full impact of pangenome-aware DV since in the GRCh38 reference out of 63.4 Mb of SDs with identity greater than 99% only 12.2 Mb was present in the T2T-Q100 high-confidence bed file. In the future, with expanding the truth set to contain more of the difficult regions we can train and test the pangenome-aware models more comprehensively.

We also counted the number of FP and FN calls induced after using pangenome-aware DV. The total number of induced errors genome-wide were 5,497 which is nearly half of the total number of fixed errors (9,272) (**Supplementary Table 14**). After computing the rate of inducing errors in the 4 partitions mentioned above we found that “SDs with identity greater than 99%” had the highest rate (**Supplementary Tables 16 and 17).** We hypothesize that this is due to the model’s reliance on pangenome information in these highly repetitive regions where short read mapping is difficult.

We also analyzed if the presence of specific allele in the population with higher frequency makes the model more confident during variant calling (**Supplementary Figures 5, 6, 7 and 8**). In order to measure that, we took the variants fixed by pagenome-aware DV and extracted their pangenome allele frequencies from the HPRC-v1.1 raw vcf file. Majority (62.8%) of removed FPs had pangenome allele frequency lower than 20% however for rescued FNs 46.3% had such low frequency (**Figure 3f**). We also extracted the pangenome allele frequencies for FPs and FNs that were induced after using pangenome-aware DV. This time the percentages of the induced FPs and FNs that had low pangenome allele frequency (lower than 20%) were 54.9% and 66.5% respectively, suggesting that having low allele frequency can be misleading for rescuing variants (**Figure 3g**; **Supplementary Figures 9, 10, 11 and 12**). As the pangenomes become more comprehensive, we expect further improvements in variant calling accuracy.

## Discussion

In this work we present pangenome-aware DeepVariant, a read-mapping-based variant caller that incorporates pangenome information to achieve high accuracy in calling sample-specific variants by augmenting input read-pileups with pangenome haplotypes. We show that pangenome-aware DeepVariant improves variant calling in different parts of the genome, with the greatest error reduction in segmental duplications with sequence identity greater than 99%. In those regions, short reads are prone to mismapping due to sparsity of unique mapping markers. We believe that the pangenome information provides additional unique markers as context, helping the model to disambiguate true variants in regions with many incorrectly mapped reads.

At present, the current T2T-Q100-v1.1 benchmark is still missing ∼80% of the segmental duplications with identity greater than 99%. Extending the truth set through collaborations between Telomere-to-Telomere and Genome-In-A-Bottle consortia remains critical as it will improve models by expanding training examples of difficult regions, and to help assess variant calling performance across samples of many ancestries.

To generalize across samples and sequencing technologies, we train with both Illumina and Element sequencing data on six different samples and three truth sets (Genome in a Bottle v4.2.1, Platinum Pedigree, and HG002 T2T-Q100). The inclusion of non-Illumina data is important for enabling users of all sequencing platforms to benefit from pangenome advancements. We anticipate pangenome-aware models incorporating other short read technologies, including Roche SBX and Complete Genomics available in the near future.

We also explored using personalized pangenomes with pangenome-aware DeepVariant, showing comparable results whether using a personalized pangenome or a full pangenome. This functionality will be critical as the pangenome size scales from the 88 haplotypes in the first release to the anticipated HPRC releases containing 1000 haplotypes^28^, ensuring DeepVariant will be compatible with and continue to benefit from the increased pangenome scale.

We further showed accuracy considerations for pangenome sampling beyond the scalability concerns. Specifically, we observed that though pangenomes could fix ∼9.3k SNP errors; however it also induced ∼5.5k SNP errors genome wide, noting that FP calls were induced because of having high frequency in the HPRC panel. This homogenizing effect would disproportionately add errors to the results for people who had ancestors living in larger populations over evolutionary time, such as existed in Africa, because larger populations are better able to retain alleles^29^. Combining personalized pangenomes with larger pangenome references can fix this issue by intelligently sampling to the most informative haplotypes that do not contain the false allele. As a result, we anticipate future HPRC graphs will improve performance further.

Methodologically, there are several approaches which can improve pangenome sampling further. The current haplotype sampling available through vg toolkit models recombination events by splitting the genome into chunks of size 10kb and then using kmers from short reads as a proxy to measure similarity between the haplotype chunks and the sample’s genome^22^.

More sophisticated approaches to splitting and indexing the pangenome for retrieval could improve performance, for example by modelling recombination hotspots^30,31^, or by going beyond k-mer approaches for measuring similarity in highly repetitive regions.

In regards to performance improvement, recent advancements in sequencing technologies and bioinformatics methods can create high-quality telomere-to-telomere (T2T) assemblies. This has enabled researchers to create pangenome references for various species and populations. In this study we showed one example application of improving an analysis method using the valuable prior information about a population. We anticipate further improvements in T2T genome assembly to enhance the usefulness in such applications. Accuracy improvements in T2T quality to fix the remaining types of errors^12^ present in pangenome assemblies will improve variant calling accuracy. Scalability of T2T assemblies will enable affordable non-human pangenomes with a quality suitable for use in non-human variant detection. Although our approach can easily be adapted for non-human species, there is a further need to generate training datasets for other species. Overall, the combination of the increasing capability of T2T assembly and pangenome creation with the ability to intelligently sample pangenome contents points towards a future with widespread, highly accurate use of population resources in genome analysis.

## Methods

### HPRC pangenome graph and GBZ format

The Human Pangenome Reference Consortium (HPRC) selected 44 samples from genetically diverse backgrounds for their first human pangenome reference. They generated high-quality phased diploid assemblies using the Trio-Hifiasm assembler with PacBio High Fidelity (HiFi) and parental Illumina short reads. Finally they constructed a pangenome graph with Minigraph-Cactus using the 88 assembled haplotypes^32^.

The HPRC pangenome is available in different formats; we selected the GBZ format for integrating with pangenome-aware DV. GBZ is a binary format primarily based on data structures from the giraffe short read mapper, and it uses GBWT index for compressing haplotype paths within the graph ^33^. Giraffe mapper and the haplotype sampling program in vg toolkit can use this format directly. Using GBZ as the input pangenome format for pangenome-aware DV eliminates the need for converting between different graph formats and enables future users to use a single pangenome file for creating pangenome-assisted read mappings, creating personalized pangenomes, and performing variant calling with pangenome-aware DV. The gbwtgraph C++ library provided an API for performing fast query operations on GBZ ^34^ and we used this API for loading GBZ and parsing haplotype alignments in pangenome-aware DeepVariant.

The pangenome resources for the first HPRC release is available in their git repository (with the commit version we used for our work); https://github.com/human-pangenomics/hpp_pangenome_resources/tree/74553c422af5521e3d297d6511da0a33fcf3a744

We used the GRCh38-based HPRC-v1.1 graph available in the GBZ format in this link: https://s3-us-west-2.amazonaws.com/human-pangenomics/pangenomes/freeze/freeze1/minigraph-cactus/hprc-v1.1-mc-grch38/hprc-v1.1-mc-grch38.gbz

### Reducing pangenome memory usage

Pangenome-aware DeepVariant can be run in a multi-threaded mode which needs splitting the genome into multiple shards and running one DeepVariant process per shard. In our initial implementation we loaded one instance of the whole graph from GBZ into memory per shard and it consumed a large size of memory (∼8 GB per shard). This implementation could not be scaled easily and needed much larger machines than the original DeepVariant. For example when we executed pangenome-aware DV with 96 threads it took ∼800Gb of memory.

To reduce pangenome memory usage we moved two memory-consuming objects into shared memory. The two objects we selected for this aim were

- std::vector<char, CharAllocatorType>* strings
- int_vector<> index

Both of which were defined inside the StringArray class of the gbwt library which is imported internally by the higher-level gbwtgraph library. These two objects were taking about ∼6.6 Gb per shard so by keeping them inside a shared memory segment we could reduce the pangenome memory usage by ∼80%. For constructing, reading, writing and destructing shared memory segments we used the interprocess API of the Boost C++ library ^35^. This API enables saving the graph objects into a shared memory segment and then mapping the segment to the address space of each DV process. Therefore each DV process could have access to the loaded graph and send queries in parallel with other processes for retrieving the haplotype alignments.

The modified library is available in this github fork: https://github.com/mobinasri/gbwtgraph

### Creating pileup images augmented with pangenome

Pangenome-aware DV takes the sample reads mapped to a reference and saved in the BAM format. It looks for candidate variants by iterating over read alignments and finding the reference positions with at least two reads supporting the same mismatch or indel. For each candidate, Pangenome-aware DV generates an image of the overlapping reads with a width of 221 bases centered on the candidate position. The height of the pileup image is set to 100 by default. This image contains multiple mandatory and optional channels, each one representing a specific type of information such as read bases, base qualities, mapping qualities, candidate support by the mappings, etc (**Supplementary Figure 1**). These read pileup images are then passed to a Convolutional Neural Network (CNN) so it can report genotype likelihoods of the candidate variants and identify false positives.

In order to make the CNN model aware of the pangenome haplotypes we added pangenome information into the input pileup images. For this aim we repurposed the DeepSomatic^36^ codebase in which the read mappings related to tumor and normal samples are layered on top of each other so that the height of the image is twice as large as linear-reference-based DeepVariant (2 x 100). Using this strategy for pangenome-aware DV we encoded the same channels for pangenome haplotypes and layered them on top of read pileups. Some of the channels like read base quality, mapping quality, and strand orientation are left blank or unused for pangenome haplotypes since they were not defined in the pangenome graph or not provided by gbwtgraph API.

Since the HPRC-v1.1 pangenome contains 88 haplotypes, using the default height of 100 was sufficient for illustrating all the haplotypes in the image. However for future releases of the HPRC pangenomes with higher number of haplotypes this height will not be enough. In that case we can increase the image height however that leads to longer runtimes for both training and testing with pangenome-aware DV since size of the CNN model needs to be scaled proportionally. The alternative approach is haplotype sampling and creating personalized pangenomes with fewer haplotypes. This approach is favored due to decreased runtime and besides that, as shown in the results, the performance with a personalized pangenome is comparable (benchmarked against T2T-Q100, **Figure 2a**) to the full pangenome.

### Truth sets and training dataset

We used three different truth sets for training pangenome-aware DV. The first one was created by the Genome-In-A-Bottle (GIAB) consortium and is one of the commonly used truth sets for benchmarking and optimizing variant calling pipelines. This truth set is available for 7 samples including Ashkenazi and Chinese trios (HG001 to HG007) however we used only 4 of them (HG004 to HG007) for training. HG003 was excluded to be used later for testing and for training with HG001 and HG002 we used other truth sets. The GIAB-v4.2.1 truth sets for HG004 to HG007 samples covers 2.51 Gb of the GRCh38 reference with 3,740,604 small variants on average.

Although GIAB-v4.2.1 has been invaluable for assessing and developing variant callers, it has limitations. One being that it is highly dependent on Illumina data in homopolymer runs and tandem repeats. The other drawback is being conservative and not covering ∼18.4% of the GRCh38 reference (for the HG003 sample).This excludes the most difficult parts of the genome for variant calling, making the assessments biased toward easy-to-call regions and not fully reflecting improvements in other parts. In order to avoid being restricted by these limitations we trained DeepVariant using two additional truth sets with more comprehensive call sets; T2T-Q100-v1.1^24,25^ for HG002 and Platinum Pedigree^37^ for HG001.

For the HG002 sample we used a more complete and accurate truth set, which was the output of the collaboration between the Genome-In-A-Bottle and the Telomere-to-Telomere (T2T) Consortia. This sample was the first diploid sample that the T2T consortium aimed to assemble in a haplotype-aware manner from telomere to telomere without any error. They had multiple iterations of assembling with the Verkko assembler using HiFi and ONT reads, filling the gaps and polishing small errors both with automated pipelines and manual investigation ^38^. The final diploid assembly is referred to as HG002-T2T-Q100 (named Q100 because it refers to the target of this effort to reach the Phred quality score of 1 error per 10 billion bases). It was then employed by the GIAB consortium for mapping to GRCh38 and creating a new truth set for HG002 in the coordinates of GRCh38. We used this truth set for training with HG002 covering 2.73 Gb of GRCh38 with 4,521,735 small variants, containing 16% more variants and spanning ∼6.8% more of the genome compared to its equivalent GIAB-v4.2.1 truth set.

For the HG001 sample we used the Platinum truth set created with a pedigree-based approach^37^. This truth set was created with read-alignment-based variant callers and using reads from various different sequencing platforms such as Illumina, PacBio HiFi and Oxford Nanopore Technologies. By sequencing the members of a large pedigree related to HG001 they were able to filter the calls violating Mendelian inheritance. Similar to GIAB-v4.2.1, Illumina was the key short read platform for creating this truth set, as evidenced by the fact that 21% of the indels in this truth set were supported only by Illumina reads and not by any other platform ^37^ . Compared to GIAB-v4.2.1 it is more comprehensive, containing 23% more variants and spanning 9.2% more of the GRCh38 reference.

Using a combination of truth sets; GIAB-v4.2.1 for HG004-HG007, T2T-Q100 for HG002 and Platinum for HG001 we made a new training dataset with the aim of having a single model that can leverage the different levels of completeness and accuracy in the available truth sets.

For training short-read pangenome-aware DeepVariant models we used the BAM files from the training dataset created for the linear-reference-based DeepVariant (https://github.com/google/deepvariant/blob/r1.8/docs/deepvariant-details-training-data.md). However we excluded all the BAM files that were mapped to the GRCh37 reference since in the HPRC-v1.1 graph GRCh37 was not present as a reference so that we could not extract the haplotype mappings to this reference. In our training dataset we had 44 BAM files with 13 and 31 of them containing Element and Illumina short reads. (**Supplementary Table 1**)

The truth set vcf files and the related high-confidence bed files are listed in the methods section, Data Sources.

### Creating personalized pangenomes

One of the major benefits of having a pangenome for mapping and variant calling happens when the variation in the sample also exists in the pangenome. That will guide the pangenome-aware mappers to locate reads properly and pangenome-aware variant callers to rescue false negatives. However, having variants that exist in the pangenome but not in the sample will be misleading and may induce read mismappings or false positive variants. Therefore removing the haplotypes that are highly divergent from the sample’s genome can potentially improve the performance of pangenome-aware DV, and is unlikely to reduce it even while allowing the number of haplotypes presented to the model to be reduced. The vg toolkit provides a haplotype sampling program that takes the kmers extracted from sample-specific reads and uses those kmers as a proxy to score the pangenome haplotypes based on their similarity to the sample’s genome. More specifically, it splits haplotypes into segments of size 10kb and scores each segment of the haplotype separately. This segmentation allows the tool to consider recombination events to some extent; however, it does not account for the case where a recombination happens within a segment. It will then output the requested number of haplotypes with the highest similarity to the sample’s genome.

We used the haplotype sampling workflow available in the vg toolkit git repo to create haplotype sampled graphs for all our training and test short read BAM files https://github.com/vgteam/vg_wdl/blob/master/workflows/haplotype_sampling.wdl

More details about the workflows we used: https://github.com/mobinasri/research_notes/blob/main/DeepVariant_Pangenome/Prepare_workflows_and_tables_for_haplotype_sampling/README.md

The pangenome-aware models were trained and tested using personalized pangenomes with 16, 32, and 64 haplotypes. 32 haplotypes were chosen to balance the tradeoff between program runtime and the accuracy of the output call set. More haplotypes lead to a taller input pileup image and a larger CNN model, which increases total inference runtime.

### Variant calling with GATK

The germline variant calling workflow with GATK ^39,40^ was written in Workflow Description Language (WDL) and is available in the WARP GitHub repository https://github.com/broadinstitute/warp/blob/develop/pipelines/broad/dna_seq/germline/single_sample/wgs/WholeGenomeGermlineSingleSample.wdl. By adjusting the parameters of the workflow, GATK can be executed in three modes. Two of these modes incorporate the submodules initially developed for the DRAGEN variant caller, aimed at enhancing indel calling in short tandem repeats and detecting correlated pileup errors ^41,42^. The 3 modes of GATK are listed below.

1. **Regular GATK (without DRAGEN modules):** It uses default values for two different versions; v3 and v4.6.
2. **GATK-DRAGEN (functional equivalence mode):** (using v4.6) By setting the parameter “dragen_functional_equivalence_mode = true”. In this mode the output of the pipeline is functionally equivalent to the one produced with the DRAGEN hardware.
3. **GATK-DRAGEN (maximum quality mode):** (using v4.6) By setting the parameter “dragen_maximum_quality_mode = true”.

In this mode the pipeline uses DRAGEN mapper and some of its variant calling steps to obtain the most accurate call set, however the final results are not functionally equivalent to the one produced with the DRAGEN hardware.

These two versions of regular GATK (v3 and v4.6) showed comparable performance while being benchmarked against GIAB-v4.2.1 and T2T-Q100-v1.1. The two modes of GATK-DRAGEN, functional equivalence and maximum quality, while being roughly comparable to each other, were significantly better than regular GATK by having about 50% fewer total errors. (**Supplementary Tables 2 and 4**)

More details about running GATK for this study is available here: https://github.com/mobinasri/research_notes/blob/main/DeepVariant_Pangenome/External_Variant_Callers/GATK/README.md

### Creating BWA and samtools indices for GRCh38 fasta

For mapping Illumina and Element short reads to GRCh38 with bwa-mem we had to create indexes with bwa and samtools.

\# bwa index

bwa index GCA_000001405.15_GRCh38_no_alt_analysis_set.fa

\# samtools index

samtools faidx GCA_000001405.15_GRCh38_no_alt_analysis_set.fa

\# list files created with bwa/samtools index ls

GCA_000001405.15_GRCh38_no_alt_analysis_set.fa

GCA_000001405.15_GRCh38_no_alt_analysis_set.fa.bwt

GCA_000001405.15_GRCh38_no_alt_analysis_set.fa.sa

GCA_000001405.15_GRCh38_no_alt_analysis_set.fa.amb

GCA_000001405.15_GRCh38_no_alt_analysis_set.fa.fai

GCA_000001405.15_GRCh38_no_alt_analysis_set.fa.ann

GCA_000001405.15_GRCh38_no_alt_analysis_set.fa.pac

### Mapping short reads with bwa mem

Here is the command we used for mapping Illumina and Element short reads with BWA-MEM after creating indexes for GRCh38. They were used for assessing Octopus and different modes of DeepVariant on BWA-MEM mapping. For running GATK and DRAGEN we didn’t use these bam files since they had their own mapping workflow.

\# for mapping Element we used the same -R value bwa mem -t 32 \

-R “@RG\tID:0\tSM:HG003.novaseq\tLB:Broad\tPU:Illumina” \

${GRCH38_FASTA} \

${FASTQ_R1} ${FASTQ_R2} | \

samtools view -bh | \

samtools sort -@ 4 -o ${OUTPUT_BAM}

\# index bam

samtools index ${OUTPUT_BAM}

### Mapping short reads with vg giraffe

We ran these commands to map short reads with vg giraffe:

- Put the paths names into a file:

cat > ${SAMPLE_NAME}.fq.paths <<- EOM

${PATH_TO_FASTQ_R1}

${PATH_TO_FASTQ_R2}

EOM

- Run kmc to create a kmer database from short reads. It will create a kff file. Temporary files will be located in a temporary directory.

TMPDIR=$(mktemp -d) kmc \

-k29 \

-m128 \

-okff \

-t$(nproc) \ @${SAMPLE_NAME}.fq.paths \

${SAMPLE_NAME}.fq \

$TMPDIR

- List the path names for GRCh38

vg paths \

-x ${PANGENOME_DIRECTORY}/hprc-v1.1-mc-grch38.gbz \

-L -Q GRCh38 > GRCh38.path_list.txt

- Run vg giraffe (v1.55) (Note that this command uses haplotype sampling for mapping reads by default)

vg giraffe --progress \

--read-group “ID:1 LB:lib1 SM:${SAMPLE_NAME} PL:${PLATFORM} PU:unit1” \

--sample ${SAMPLE_NAME} \

-o BAM \

--ref-paths GRCh38.path_list.txt \

-P -L 3000 \

-f ${PATH_TO_FASTQ_R1} \

-f ${PATH_TO_FASTQ_R2} \

-Z ${PANGENOME_DIRECTORY}/hprc-v1.1-mc-grch38.gbz \

--kff-name ${SAMPLE_NAME}.fq.kff \

--haplotype-name ${PANGENOME_DIRECTORY}/hprc-v1.1-mc-grch38.hapl \

-t $(nproc) > ${PATH_TO_UNSORTED_BAM}

- Clean up contig names, sort alignments by coordinates and index final bam

samtools view -h ${PATH_TO_UNSORTED_BAM} | \ sed -e “s/GRCh38#0#//g” | \

samtools sort --threads 10 -m 2G -O BAM > ${PATH_TO_SORTED_BAM} samtools index -@$(nproc) ${PATH_TO_SORTED_BAM}

### Variant calling with Octopus

Octopus is a read-mapping-based variant caller that locally assembles reads to create haplotypes and then builds a tree of haplotypes that will be later pruned by using the posterior likelihoods of read-haplotype pairs. These haplotypes are used for genotyping and phasing variants. The final vcf file is filtered using a random forest classifier however the config file of this classifier was not available in their github repository therefore our Octopus results for this study were not filtered.

We ran Octopus-v0.7.4 on 4 different HG003 (and HG002) bam files created using all combinations of mapping Illumina and Element reads with vg giraffe and BWA-MEM read mappers. Here is the command for running Octopus:

docker run -it -v${INPUT_DIR}:${INPUT_DIR} dancooke/octopus:latest octopus \

-R ${GRCH38_FASTA} \

-I ${INPUT_BAM} \

-T chr1 to chrM \

--sequence-error-model PCR-FREE.NOVASEQ \

-o ${OUTPUT_VCF} \

--threads 32

### Variant calling with DRAGEN-v4.3

DRAGEN-v4.3 is the recent version of Illumina’s hardware-accelerated mapper and variant caller, which we ran using their cloud-based support on AWS. We used DRAGEN for both read mapping and variant calling. We used Illumina’s most recent Multigenome reference genome hash table. This reference was extracted into the ∼/ref/directory for our analysis. The command used to run DRAGEN was as follows:

dragen -1 ${FASTQ_R1} -2 ${FASTQ_R2} \

--RGSM ${SAMPLE_NAME} \

--RGID ${SAMPLE_NAME} \

--ref-dir ∼/ref/\

--output-file-prefix ${OUTPUT_PREFIX} \

--events-log-file dragen_events.csv \

--output-directory output \

--generate-sa-tags true \

--enable-vcf-compression true \

--enable-variant-caller true \

--enable-map-align true \

--enable-map-align-output true \

--enable-sort true \

--enable-duplicate-marking true \

--enable-bam-indexing true \

--intermediate-results-dir tmp

### Variant calling with pangenome-aware DeepVariant

mkdir -p output

mkdir -p output/intermediate_results_dir

BIN_VERSION="pangenome_aware_deepvariant-head737001992"

sudo docker pull gcr.io/deepvariant-docker/deepvariant:"${BIN_VERSION}"

sudo docker run \

-v “${PWD}/input":"/input” \

-v “${PWD}/output":"/output” \

-v “${PWD}/reference":"/reference” \

--shm-size 12gb \

gcr.io/deepvariant-docker/deepvariant:"${BIN_VERSION}” \

/opt/deepvariant/bin/run_pangenome_aware_deepvariant \

--model_type WGS \

--ref /reference/GRCh38_no_alt_analysis_set.fasta \

--reads /input/${INPUT_BAM} \

--pangenome /input/hprc-v1.1-mc-grch38.gbz \

--output_vcf /output/${SAMPLE_NAME}.output.vcf.gz \

--output_gvcf /output/${SAMPLE_NAME}.output.g.vcf.gz \

--num_shards $(nproc) \

--intermediate_results_dir /output/intermediate_results_dir

#### Running hap.py

We used hap.py for benchmarking the outputs of different variant calling tools against the 3 truth sets of GIAB-v4.2.1 for HG003, T2T-Q100 for HG002, Platinum for HG001.

HAPPY_VERSION="v0.3.12"

\# male for HG002/HG003 and female for HG001 GENDER="male"

docker run -i -v${INPUT_DIR}:${INPUT_DIR} jmcdani20/hap.py:${HAPPY_VERSION} /opt/hap.py/bin/

hap.py \

${TRUTH_VCF} \

${QUERY_VCF} \

-f ${TRUTH_CONF_BED} \

-r ${GRCH38_FASTA} \

-o ${OUTPUT_PREFIX} \

--gender ${GENDER} \

--engine=vcfeval \

--pass-only

### Data Sources

DeepVariant WGS models and Docker image

- Docker image
  - gcr.io/deepvariant-docker/deepvariant:pangenome_aware_deepvariant-head737001992
- DV model with no pangenome

- https://storage.googleapis.com/brain-genomics-public/research/pangenome_aware_dv_paper/may_2025/models/original_dv/checkpoint-227328-0.98714-1.data-00000-of-00001
- https://storage.googleapis.com/brain-genomics-public/research/pangenome_aware_dv_paper/may_2025/models/original_dv/checkpoint-227328-0.98714-1.index
- https://storage.googleapis.com/brain-genomics-public/research/pangenome_aware_dv_paper/may_2025/models/original_dv/example_info.json
- Pangenome-aware with the full pangenome (88 haplotypes)

- https://storage.googleapis.com/brain-genomics-public/research/pangenome_aware_dv_paper/may_2025/models/pangenome_aware_dv_full/checkpoint-166400-0.98779-1.data-00000-of-00001
- https://storage.googleapis.com/brain-genomics-public/research/pangenome_aware_dv_paper/may_2025/models/pangenome_aware_dv_full/checkpoint-166400-0.98779-1.index
- https://storage.googleapis.com/brain-genomics-public/research/pangenome_aware_dv_paper/may_2025/models/pangenome_aware_dv_full/example_info.json
- Pangenome-aware with personalized pangenome (16 haplotypes)

- https://storage.googleapis.com/brain-genomics-public/research/pangenome_aware_dv_paper/may_2025/models/pangenome_aware_dv_16_haps/checkpoint-153600-0.98945-1.data-00000-of-00001
- https://storage.googleapis.com/brain-genomics-public/research/pangenome_aware_dv_paper/may_2025/models/pangenome_aware_dv_16_haps/checkpoint-153600-0.98945-1.index
- https://storage.googleapis.com/brain-genomics-public/research/pangenome_aware_dv_paper/may_2025/models/pangenome_aware_dv_16_haps/example_info.json
- Pangenome-aware with personalized pangenome (32 haplotypes)

- https://storage.googleapis.com/brain-genomics-public/research/pangenome_aware_dv_paper/may_2025/models/pangenome_aware_dv_32_haps/checkpoint-179200-0.98890-1.data-00000-of-00001
- https://storage.googleapis.com/brain-genomics-public/research/pangenome_aware_dv_paper/may_2025/models/pangenome_aware_dv_32_haps/checkpoint-179200-0.98890-1.index
- https://storage.googleapis.com/brain-genomics-public/research/pangenome_aware_dv_paper/may_2025/models/pangenome_aware_dv_32_haps/example_info.json

### Truth Sets and confident BED files

- GIAB-v4.2.1 for HG003

- https://ftp-trace.ncbi.nlm.nih.gov/ReferenceSamples/giab/release/AshkenazimTrio/HG003_NA24149_father/NISTv4.2.1/GRCh38/
- T2T-Q100-v1.1 for HG002

- https://storage.googleapis.com/brain-genomics-public/research/pangenome_aware_dv_paper/jan_2025/truth/hg002_t2t_v1.1/GRCh38_HG2-T2TQ100-V1.1_smvar.benchmark.bed
- https://storage.googleapis.com/brain-genomics-public/research/pangenome_aware_dv_paper/jan_2025/truth/hg002_t2t_v1.1/GRCh38_HG2-T2TQ100-V1.1_smvar_dipcall-z2k.vcf.gz
- https://storage.googleapis.com/brain-genomics-public/research/pangenome_aware_dv_paper/jan_2025/truth/hg002_t2t_v1.1/GRCh38_HG2-T2TQ100-V1.1_smvar_dipcall-z2k.vcf.gz.tbi
- Platinum for HG001

- https://storage.googleapis.com/brain-genomics-public/research/pangenome_aware_dv_paper/jan_2025/truth/hg001_platinum_pedigree/hq_regions_final.sep2024.bed
- https://storage.googleapis.com/brain-genomics-public/research/pangenome_aware_dv_paper/jan_2025/truth/hg001_platinum_pedigree/NA12878_hq_truthset.sep2024.vcf.gz
- https://storage.googleapis.com/brain-genomics-public/research/pangenome_aware_dv_paper/jan_2025/truth/hg001_platinum_pedigree/NA12878_hq_truthset.sep2024.vcf.gz.tbi

### Illumina NovaSeq reads

- HG001

- https://storage.googleapis.com/brain-genomics-public/research/pangenome_aware_dv_paper/jan_2025/fastq/HG001.novaseq.pcr-free.35x.R1.fastq.gz
- https://storage.googleapis.com/brain-genomics-public/research/pangenome_aware_dv_paper/jan_2025/fastq/HG001.novaseq.pcr-free.35x.R2.fastq.gz
- HG002

- https://storage.googleapis.com/brain-genomics-public/research/pangenome_aware_dv_paper/jan_2025/fastq/HG002.novaseq.pcr-free.35x.R1.fastq.gz
- https://storage.googleapis.com/brain-genomics-public/research/pangenome_aware_dv_paper/jan_2025/fastq/HG002.novaseq.pcr-free.35x.R2.fastq.gz
- HG003

- https://storage.googleapis.com/brain-genomics-public/research/pangenome_aware_dv_paper/jan_2025/fastq/HG003.novaseq.pcr-free.35x.R1.fastq.gz
- https://storage.googleapis.com/brain-genomics-public/research/pangenome_aware_dv_paper/jan_2025/fastq/HG003.novaseq.pcr-free.35x.R2.fastq.gz
- Element reads
- HG001

- https://storage.googleapis.com/brain-genomics-public/research/element/cloudbreak_wgs/HG001.element.cloudbreak.1000bp_ins.R1.fastq.gz
- https://storage.googleapis.com/brain-genomics-public/research/element/cloudbreak_wgs/HG001.element.cloudbreak.1000bp_ins.R2.fastq.gz
- HG002

- https://storage.googleapis.com/brain-genomics-public/research/element/cloudbreak_wgs/HG002.element.cloudbreak.1000bp_ins.R1.fastq.gz
- https://storage.googleapis.com/brain-genomics-public/research/element/cloudbreak_wgs/HG002.element.cloudbreak.1000bp_ins.R2.fastq.gz
- HG003
- https://storage.googleapis.com/brain-genomics-public/research/element/cloudbreak_wgs/HG003.element.cloudbreak.1000bp_ins.R1.fastq.gz
- https://storage.googleapis.com/brain-genomics-public/research/element/cloudbreak_wgs/HG003.element.cloudbreak.1000bp_ins.R2.fastq.gz

### vg giraffe Alignments

- HG001 Element

- https://storage.googleapis.com/brain-genomics-public/research/pangenome_aware_dv_paper/jan_2025/vg_bam/HG001.element.cloudbreak.1000bp_ins.vg.grch38.bam
- https://storage.googleapis.com/brain-genomics-public/research/pangenome_aware_dv_paper/jan_2025/vg_bam/HG001.element.cloudbreak.1000bp_ins.vg.grch38.bam.bai
- HG002 Element

- https://storage.googleapis.com/brain-genomics-public/research/pangenome_aware_dv_paper/jan_2025/vg_bam/HG002.element.cloudbreak.1000bp_ins.vg.grch38.bam
- https://storage.googleapis.com/brain-genomics-public/research/pangenome_aware_dv_paper/jan_2025/vg_bam/HG002.element.cloudbreak.1000bp_ins.vg.grch38.bam.bai
- HG003 Element

- https://storage.googleapis.com/brain-genomics-public/research/pangenome_aware_dv_paper/jan_2025/vg_bam/HG003.element.cloudbreak.1000bp_ins.vg.grch38.bam
- https://storage.googleapis.com/brain-genomics-public/research/pangenome_aware_dv_paper/jan_2025/vg_bam/HG003.element.cloudbreak.1000bp_ins.vg.grch38.bam.bai
- HG001 NovaSeq

- https://storage.googleapis.com/deepvariant/vg-case-study/HG001.novaseq.pcr-free.35x.vg-1.55.0.bam
- https://storage.googleapis.com/deepvariant/vg-case-study/HG001.novaseq.pcr-free.35x.vg-1.55.0.bam.bai
- HG002 NovaSeq
- https://storage.googleapis.com/deepvariant/vg-case-study/HG002.novaseq.pcr-free.35x.vg-1.55.0.bam
- https://storage.googleapis.com/deepvariant/vg-case-study/HG002.novaseq.pcr-free.35x.vg-1.55.0.bam.bai
- HG003 NovaSeq

- https://storage.googleapis.com/deepvariant/vg-case-study/HG003.novaseq.pcr-free.35x.vg-1.55.0.bam
- https://storage.googleapis.com/deepvariant/vg-case-study/HG003.novaseq.pcr-free.35x.vg-1.55.0.bam.bai

### HPRC GBZ files

- https://s3-us-west-2.amazonaws.com/human-pangenomics/pangenomes/freeze/freeze1/minigraph-cactus/hprc-v1.1-mc-grch38/hprc-v1.1-mc-grch38.gbz

### DRAGEN reference hash table

- https://webdata.illumina.com/downloads/software/dragen/resource-files/misc/hg19-alt_masked.cnv.graph.hla.rna-10-r4.0-1.tar

## Acknowledgements

B.P. and UCSC personnel were in part supported by NIH grants R01HG010485, U01HG010961,U24HG010262, and U24HG011853.

## Author contributions

K.S., B.P., A.C., L.D., and D.R.W. devised the study. M.A., A.C., B.P., and K.S. drafted manuscript. M.A., P.C., K.S., J.C.M., A.K., D.E.C., and L.B. developed pangenome-aware DeepVariant. J.S., G.H., and A.M.N. developed tools for constructing pangenomes, read mapping and creating personalized pangenomes. M.A. and P.E. ran and evaluated external variant calling tools.

## Supplementary Figures

**Supplementary Figure 1:**
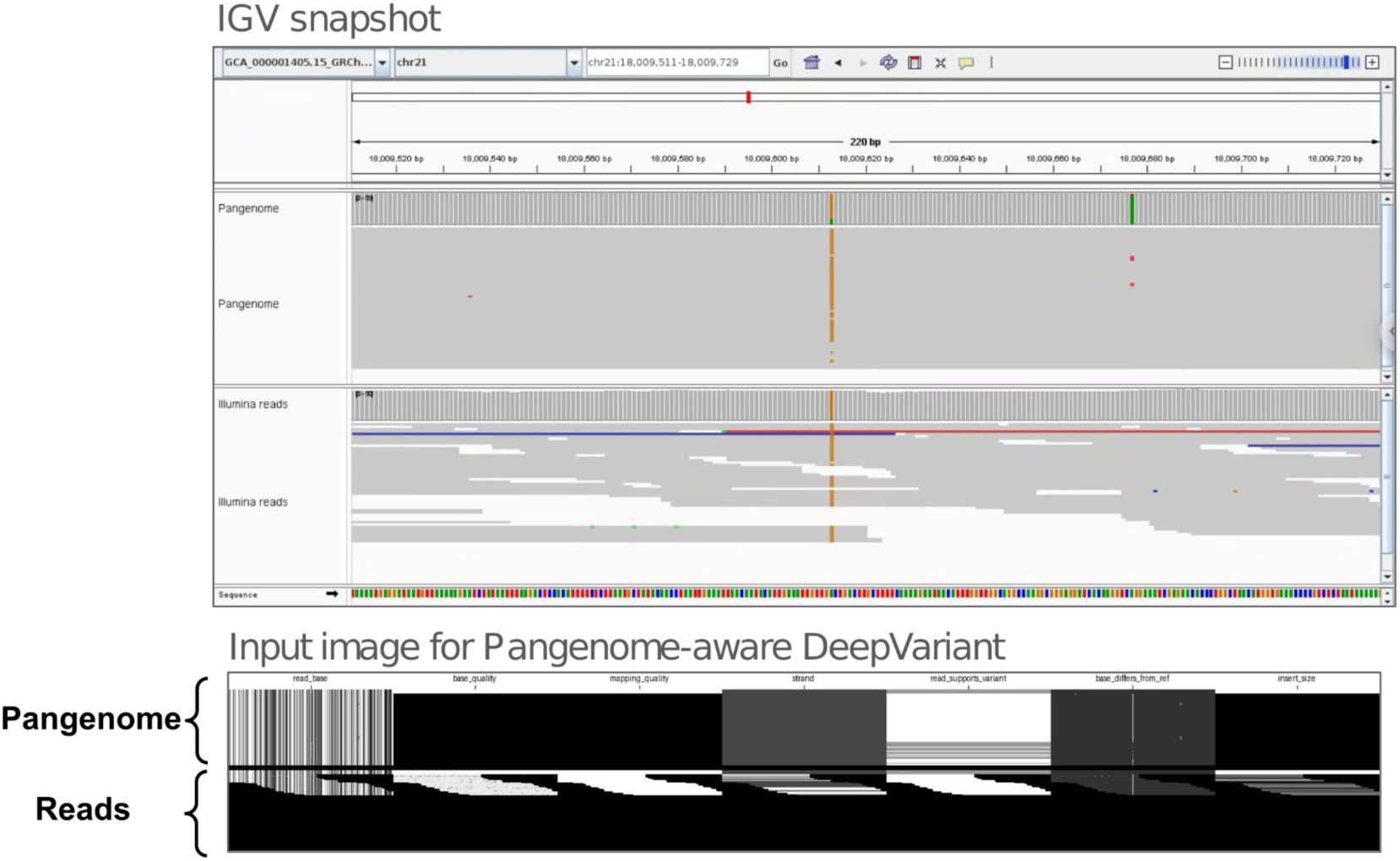
**Input pileup image for pangenome-aware DeepVariant.** The top panel shows an IGV snapshot with pangenome haplotypes and short read alignments. The bottom panel shows the related DeepVariant pileup image for the same window as the top IGV snapshot. Pangenome-aware DeepVariant creates this image and feeds it into the CNN model. The top and bottom parts of the pileup image contain pangenome haplotypes and read alignments respectively.

**Supplementary Figure 2:**
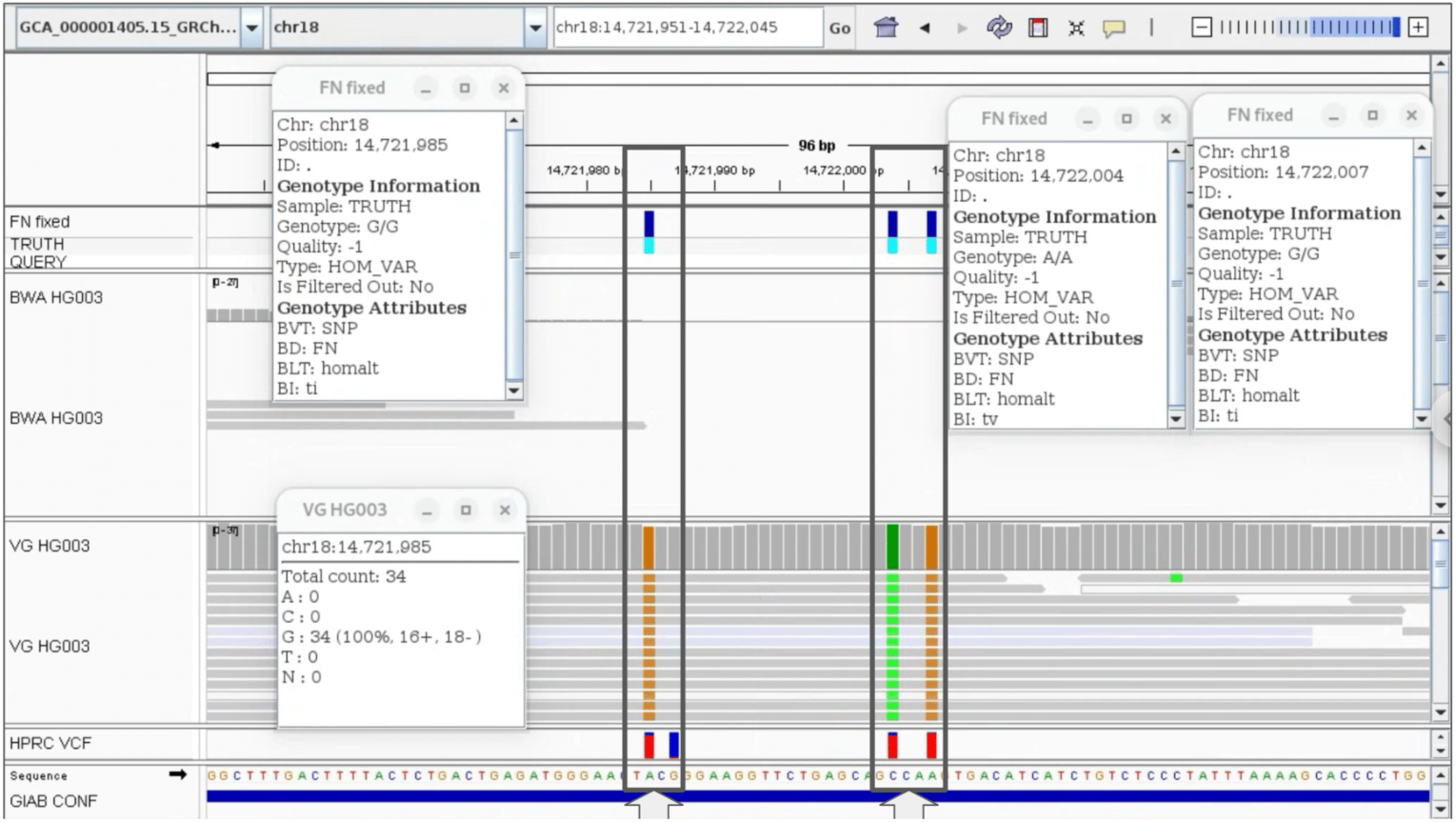
Example of false negative calls rescued by using vg giraffe mappings. linear-reference-based DV (with no pangenome) was run on HG003 NovaSeq reads mapped with vg giraffe and BWA-MEM. vg mappings provide clear signals for three true positive homozygous calls based on the GIAB-v4.2.1 truth set. This led to rescuing these calls which were missed when BWA-MEM was the mapper.

**Supplementary Figure 3:**
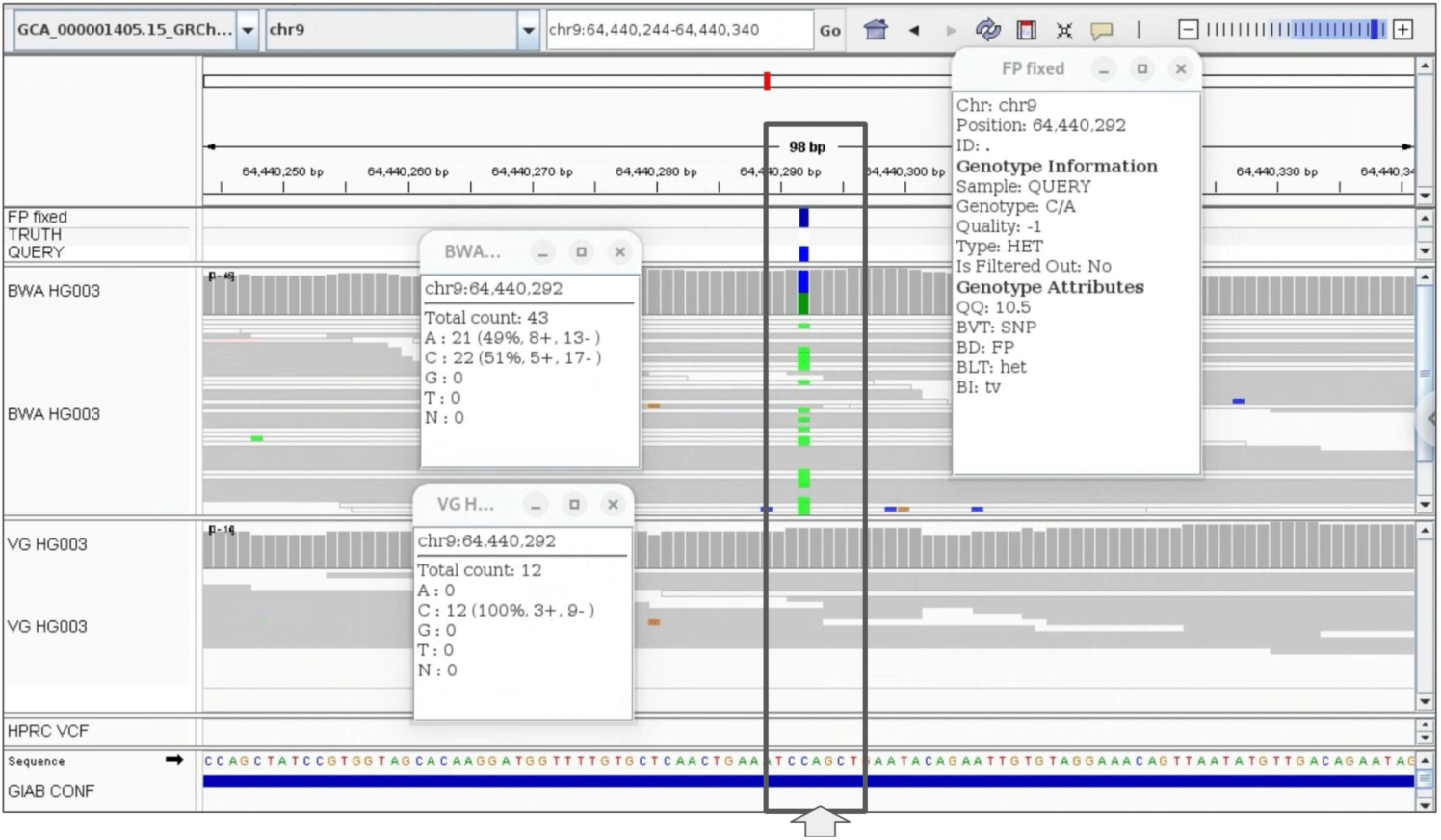
Example of a false positive call removed by using vg giraffe mappings. linear-reference-based DV (with no pangenome) was run on HG003 NovaSeq reads mapped with vg giraffe and BWA-MEM. Some of the BWA-MEM mappings contain a SNP however none of the vg mappings contains that. This led to calling a false positive heterozygous SNP with BWA-MEM, which was absent from the calls made with vg mappings.

**Supplementary Figure 4:**
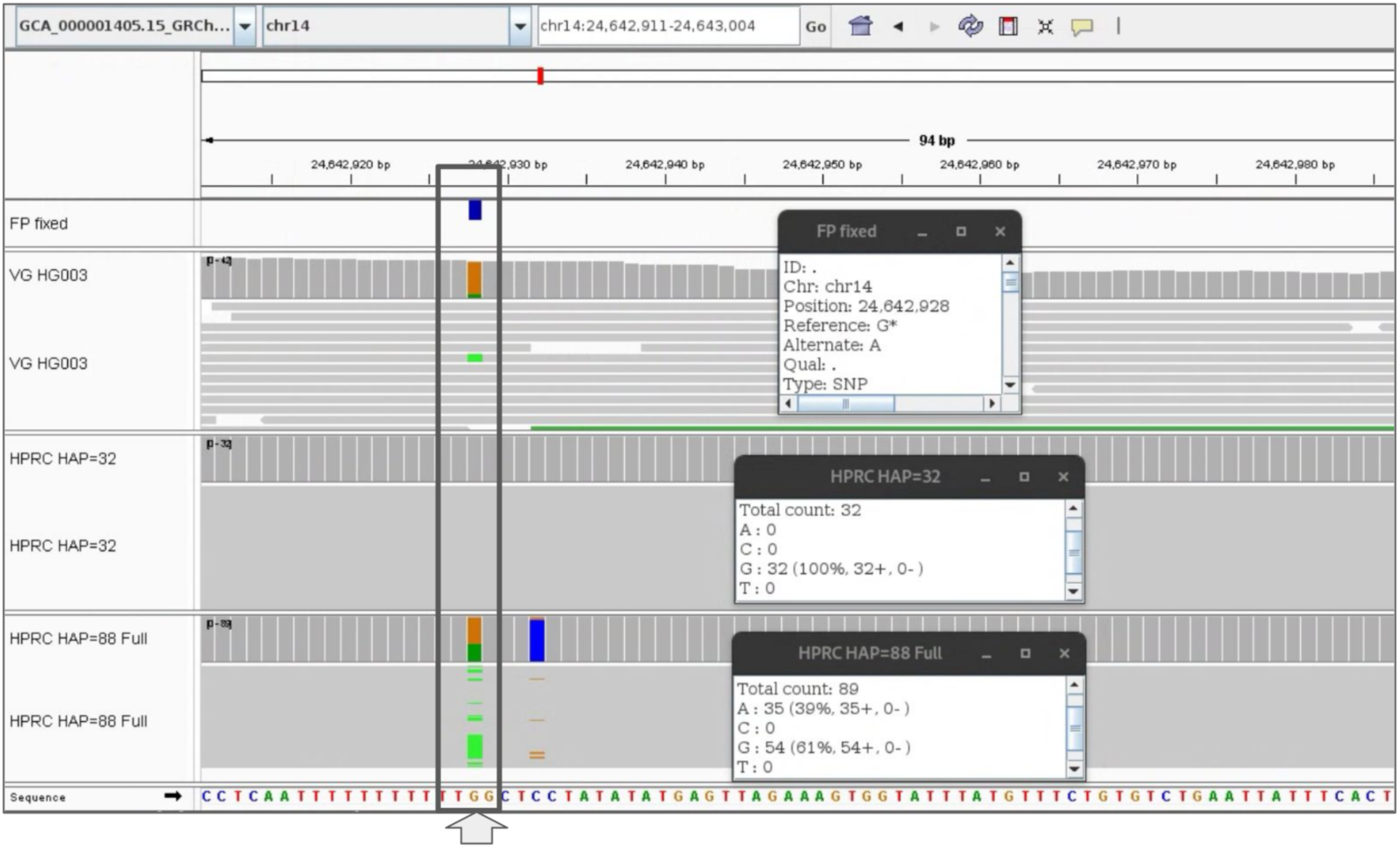
**Example of a false positive call removed by using personalized pangenome** Here is an example of how a personalized pangenome provides the haplotypes that more closely represent the sample’s genome. In this example when pangenome-aware DV was run with full pangenome (88 haplotypes + CHM13 reference) it called a false positive; however when personalized pangenome was used (with 32 haplotypes) it didn’t call it. All haplotypes with the alternative allele of “A” were removed in the personalized pangenome.

**Supplementary Figure 5:**
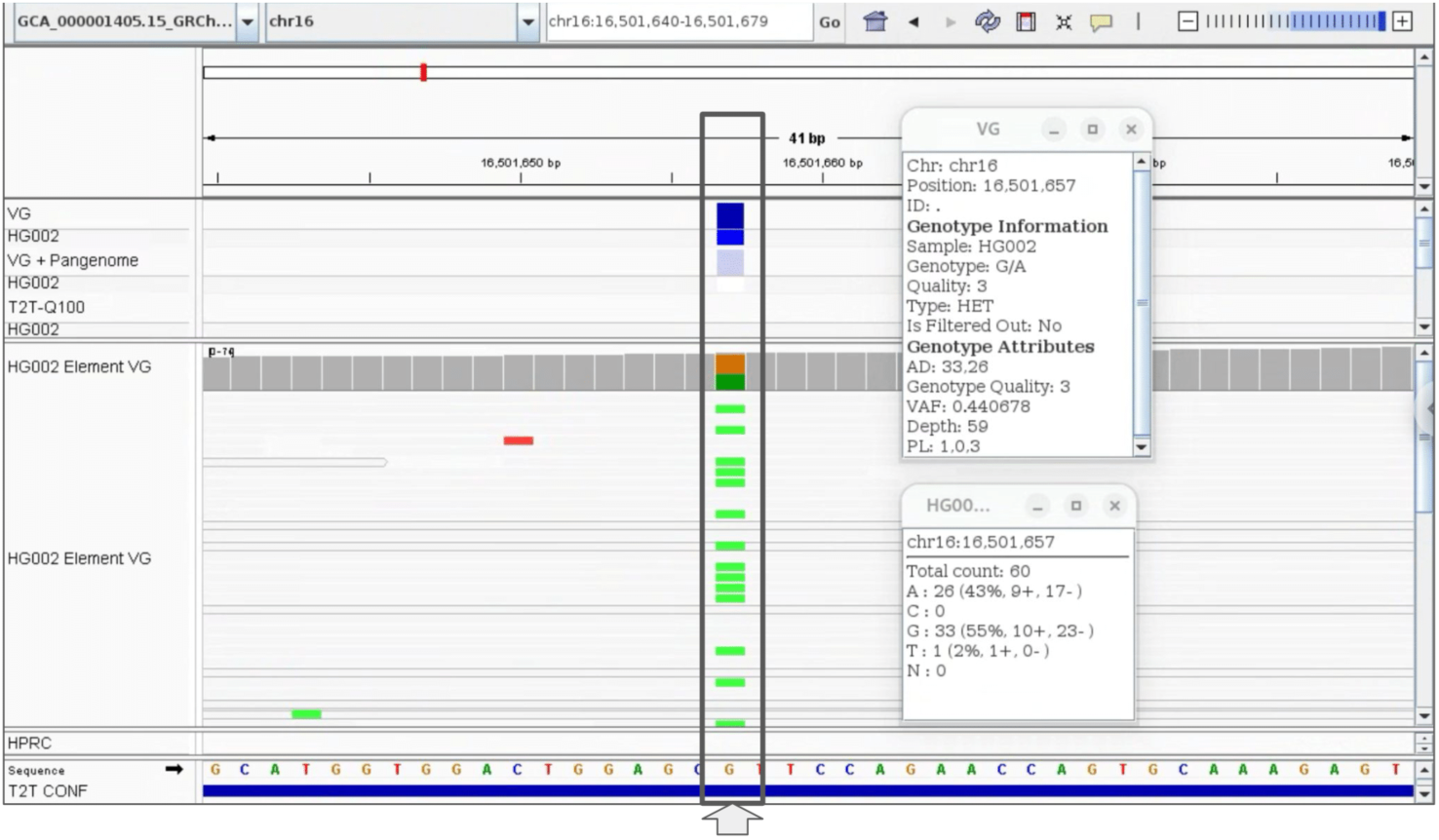
Example of a false positive call removed by pangenome-aware DV. linear-reference-based DV and pangenome-aware DV were run on HG002 Element reads mapped with vg giraffe. We show one FP call in chr16 that was removed after using pangenome-aware DV. The alternative allele was completely absent from the pangenome.

**Supplementary Figure 6:**
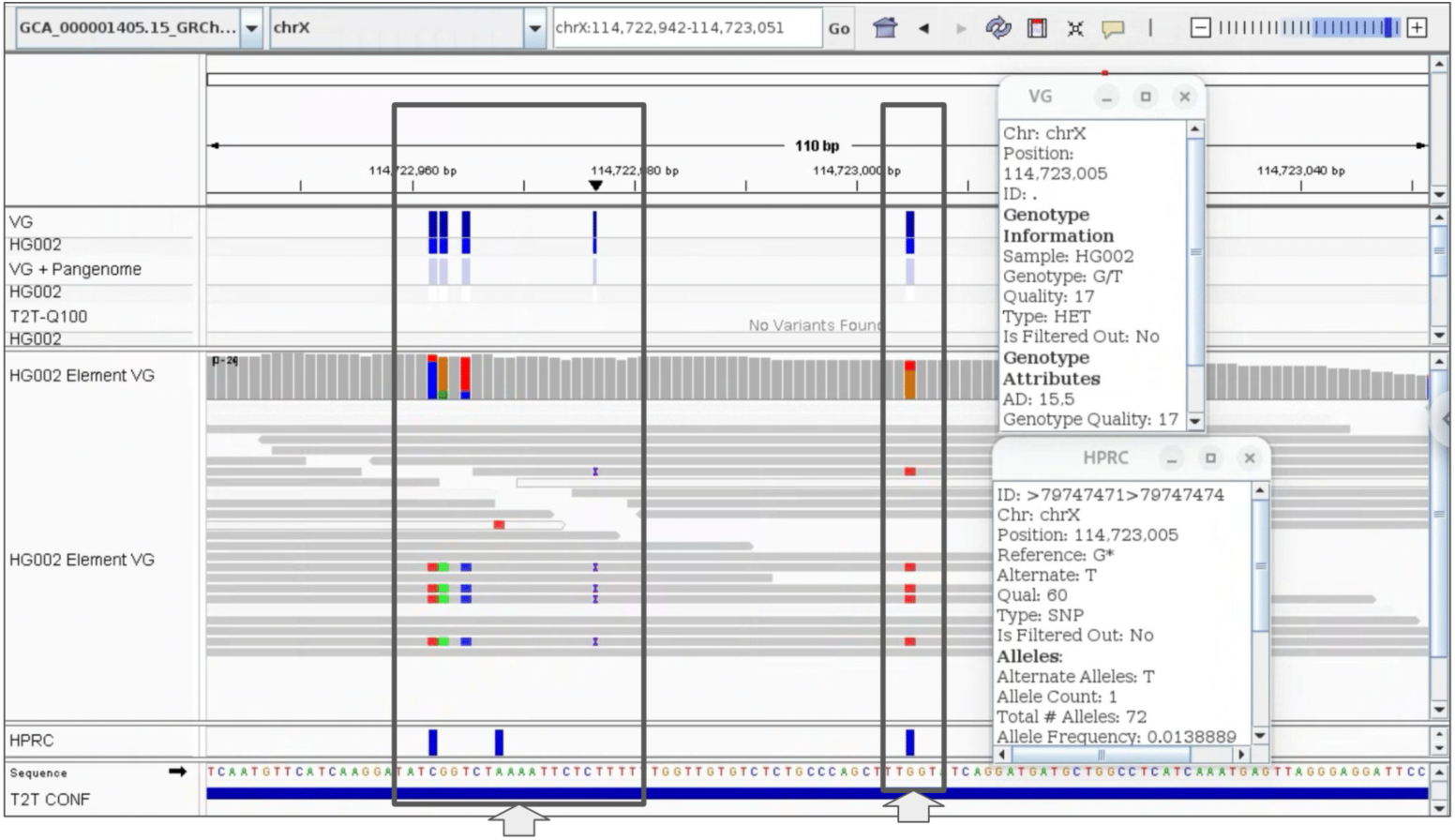
**Example of false positive calls removed by pangenome-aware DV** linear-reference-based DV and pangenome-aware DV were run on HG002 Element reads mapped with vg. We show five FP calls in chrX that were removed after using pangenome-aware DV. For example for the right-most FP SNP there is only one haplotype (out of 72) with the alternative allele in the HPRC panel.

**Supplementary Figure 7:**
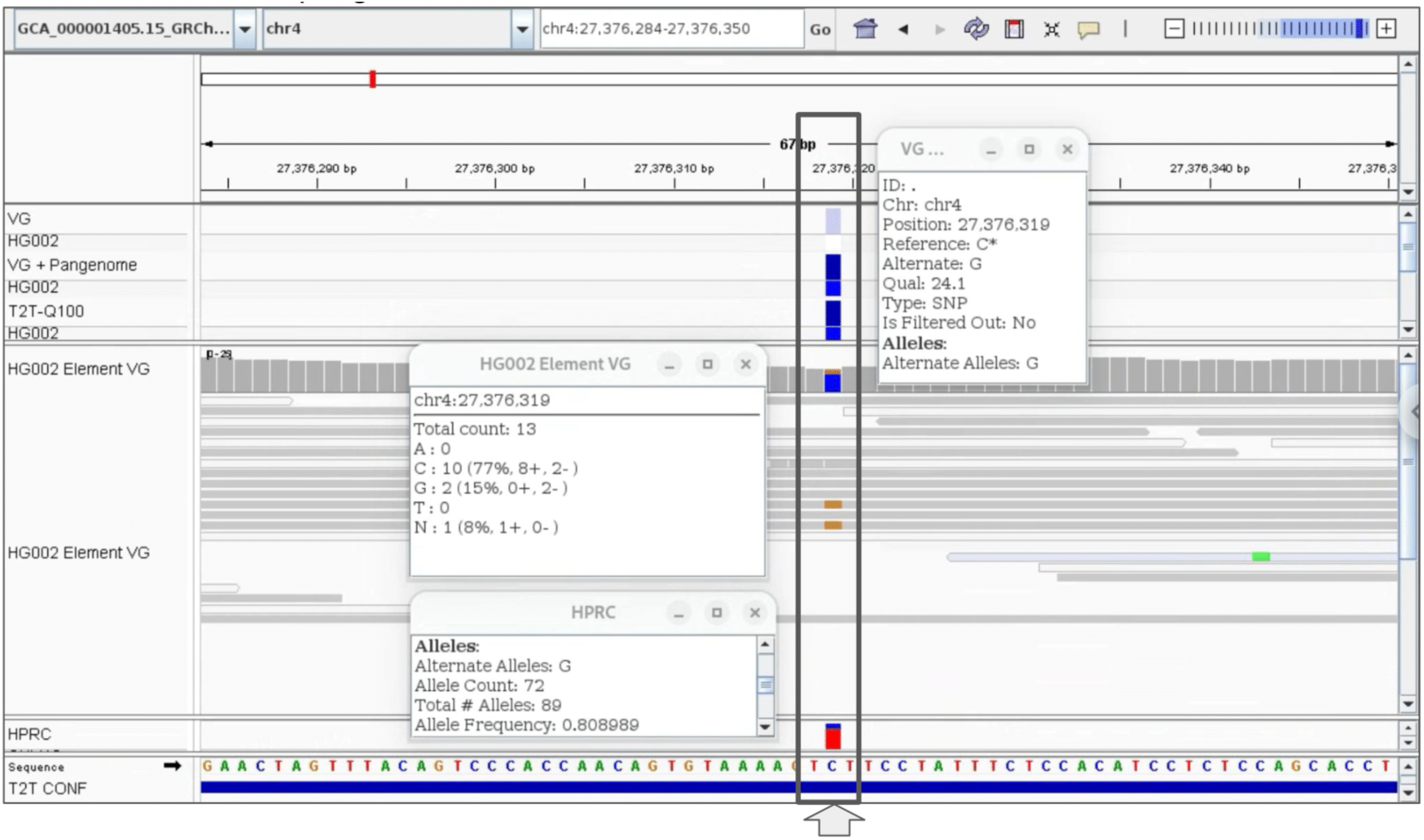
**Example of a false negative call rescued by pangenome-aware DV** linear-reference-based DV and pangenome-aware DV were run on HG002 Element reads mapped with vg giraffe. We show one FN call in chr4 that was rescued after using pangenome-aware DV. There are 72 haplotypes (out of 89) supporting this allele in the HPRC panel.

**Supplementary Figure 8:**
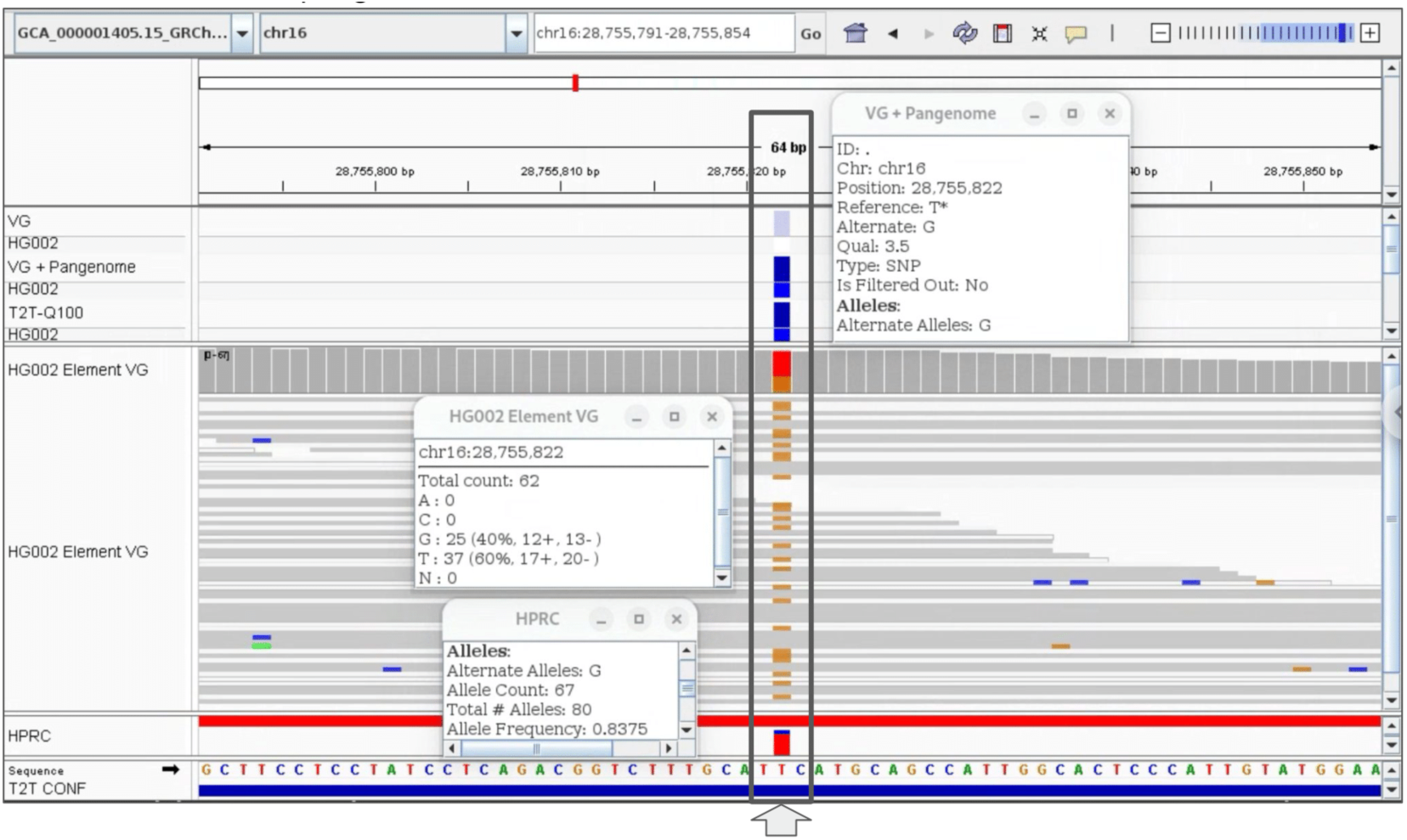
Example of a false negative call rescued by pangenome-aware DV. linear-reference-based DV and pangenome-aware DV were run on HG002 Element reads mapped with vg. We show one FN call in chr16 that was rescued after using pangenome-aware DV. There are 67 haplotypes (out of 80) supporting this allele in the HPRC panel.

**Supplementary Figure 9:**
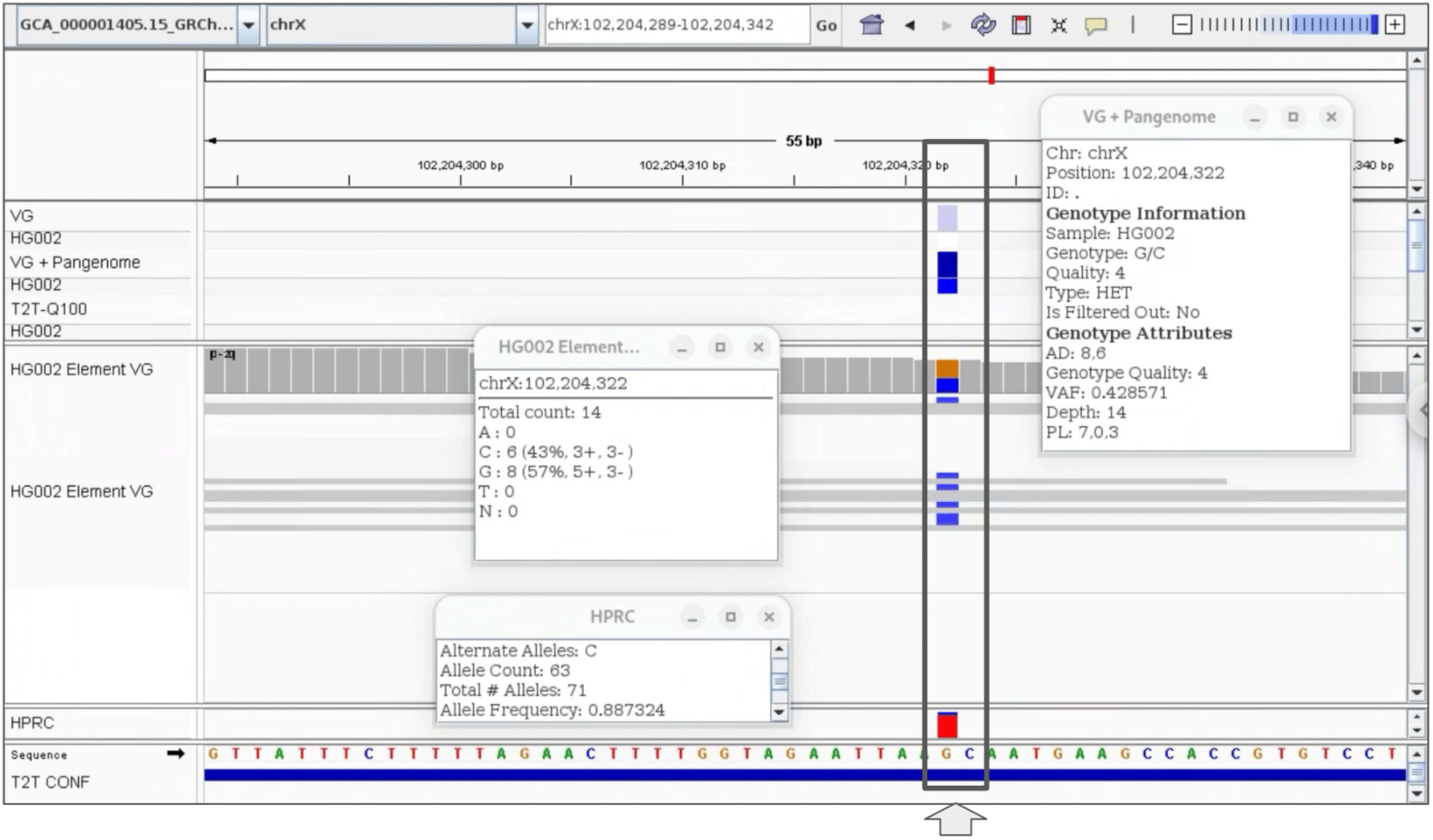
Example of a false positive call induced by pangenome-aware DV. linear-reference-based DV and pangenome-aware DV were run on HG002 Element reads mapped with vg giraffe. Pangenome-aware DV calls a FP heterozygous SNP, which was not called by the linear-reference-based DV. The alternate allele “C” is highly common in the HPRC panel (63 out of 71).

**Supplementary Figure 10:**
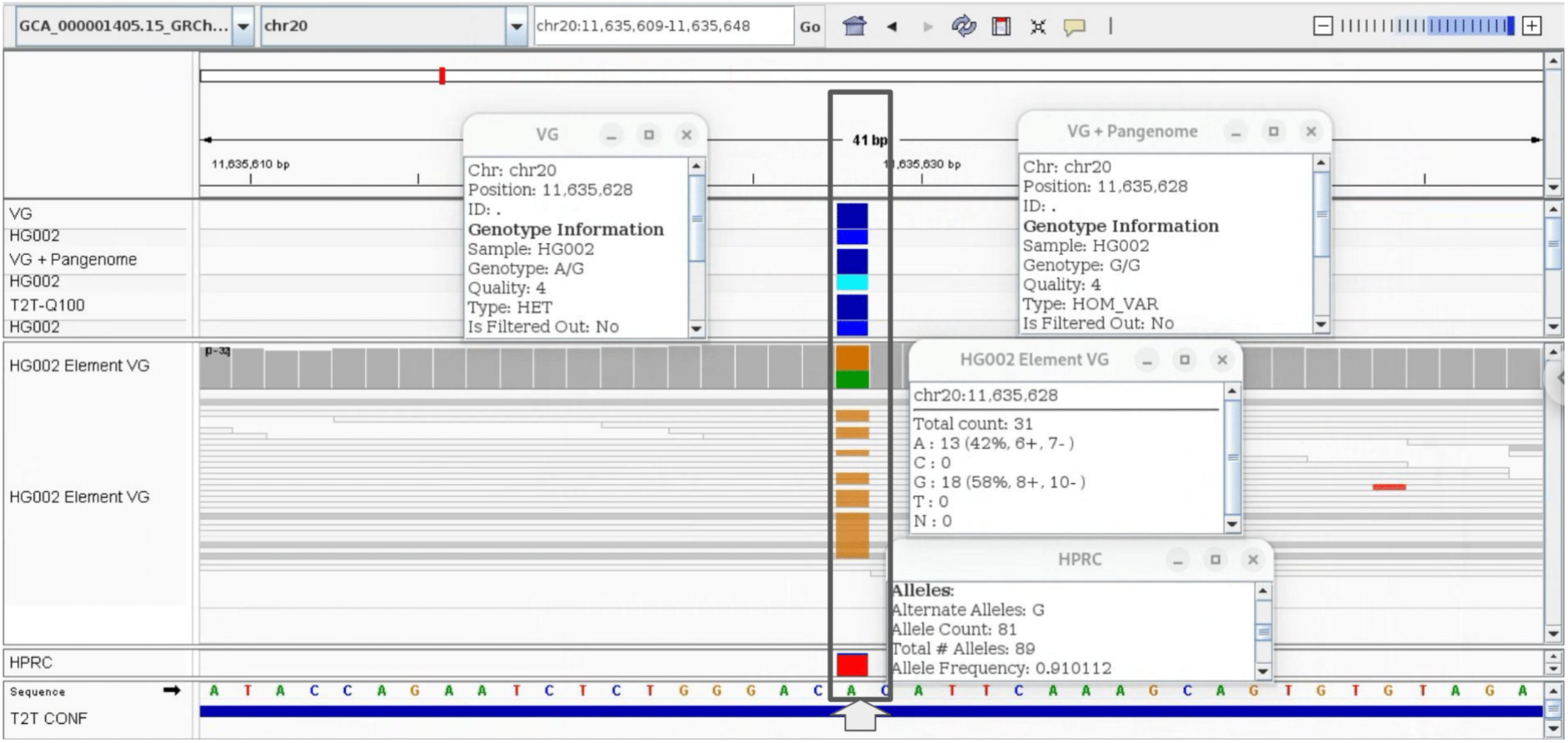
Example of a false positive call induced by pangenome-aware DV. linear-reference-based DV and pangenome-aware DV were run on HG002 Element reads mapped with vg giraffe. linear-reference-based DV calls a TP heterozygous SNP however pangenome-aware DV calls it homozygous by mistake. There are 81 haplotypes (out of 89) supporting this allele in the HPRC panel.

**Supplementary Figure 11:**
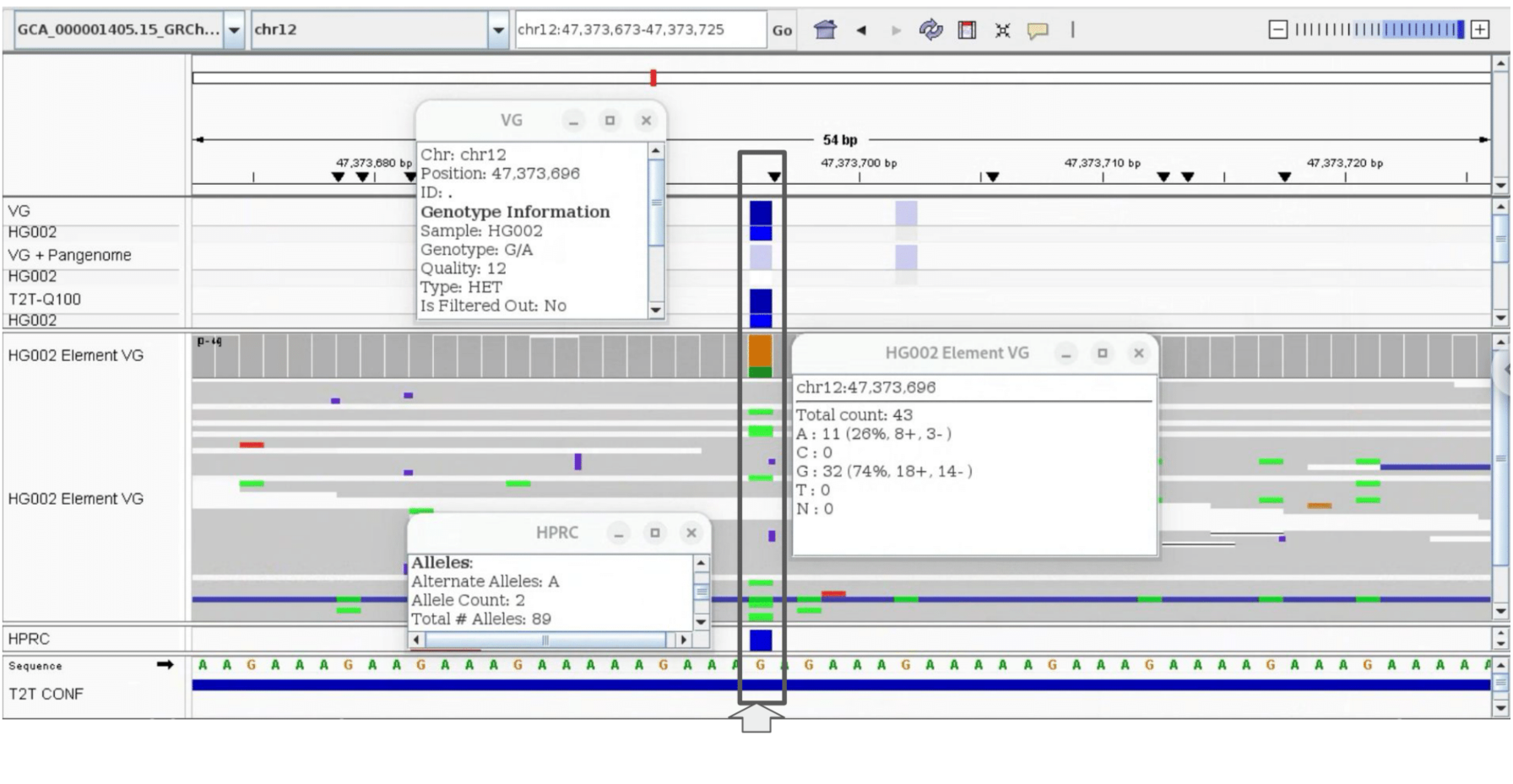
Example of a false negative call induced by pangenome-aware DV. linear-reference-based DV and pangenome-aware DV were run on HG002 Element reads mapped with vg giraffe. linear-reference-based DV calls a TP heterozygous SNP however it is missed by pangenome- aware DV. There are only 2 haplotypes (out of 89) supporting this allele in the HPRC panel.

**Supplementary Figure 12:**
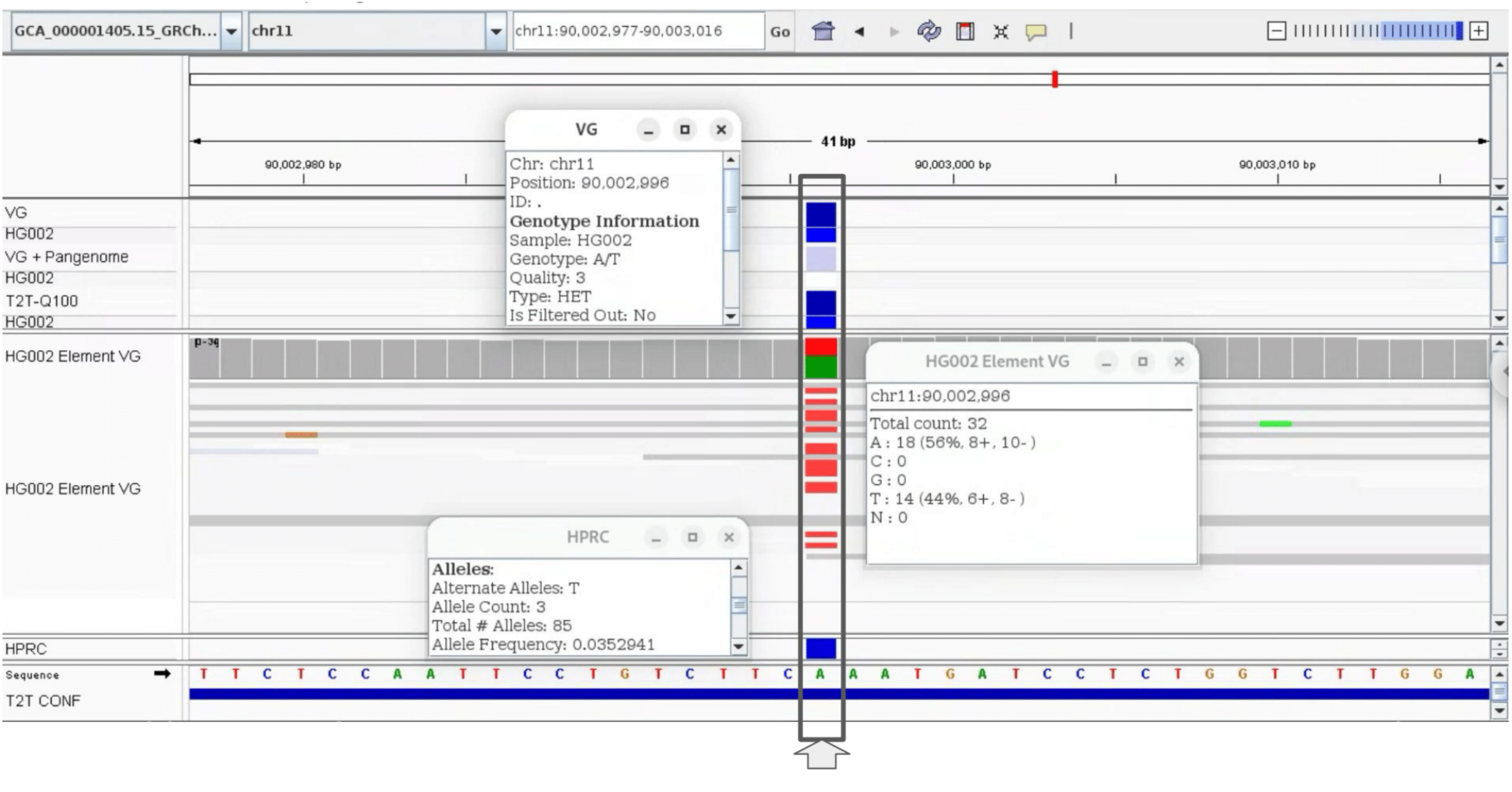
Example of a false negative call induced by pangenome-aware DV. linear-reference-based DV and pangenome-aware DV were run on HG002 Element reads mapped with vg giraffe. linear-reference-based DV calls a TP heterozygous SNP however it is missed by pangenome-aware DV. There are only 3 haplotypes (out of 85) supporting this allele in the HPRC panel.

**Supplementary Figure 13:**
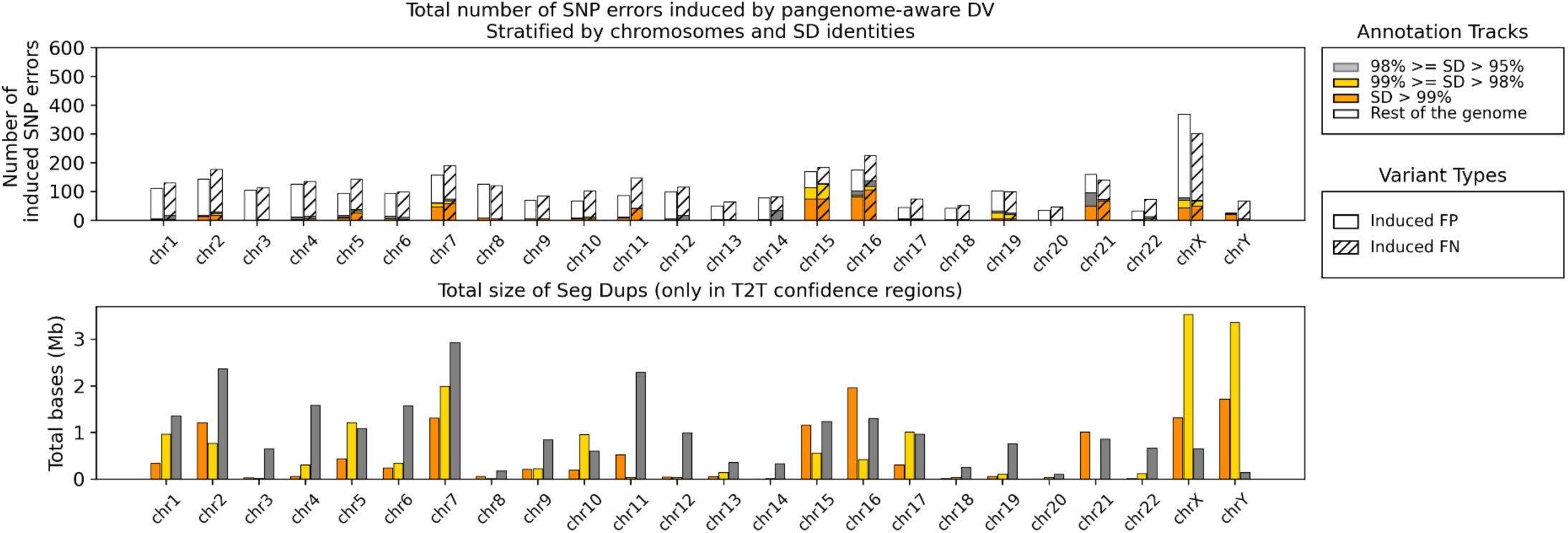
**Total number of SNP errors induced by pangenome-aware DV and total size of seg dups in T2T high-confidence regions** Top panel shows the number of SNP errors induced by pangenome-aware DV, which were absent from the linear-reference-based DV. For this analysis HG002 Element reads were mapped with vg giraffe and the T2T-Q100 truth set was used for benchmarking. This panel can be compared with Figure 3d (equivalent figure but for fixed SNP errors). The bottom panel shows the total length of segmental duplications in the high-confidence bed file stratified by identity across all chromosomes.

**Supplementary Figure 14:**
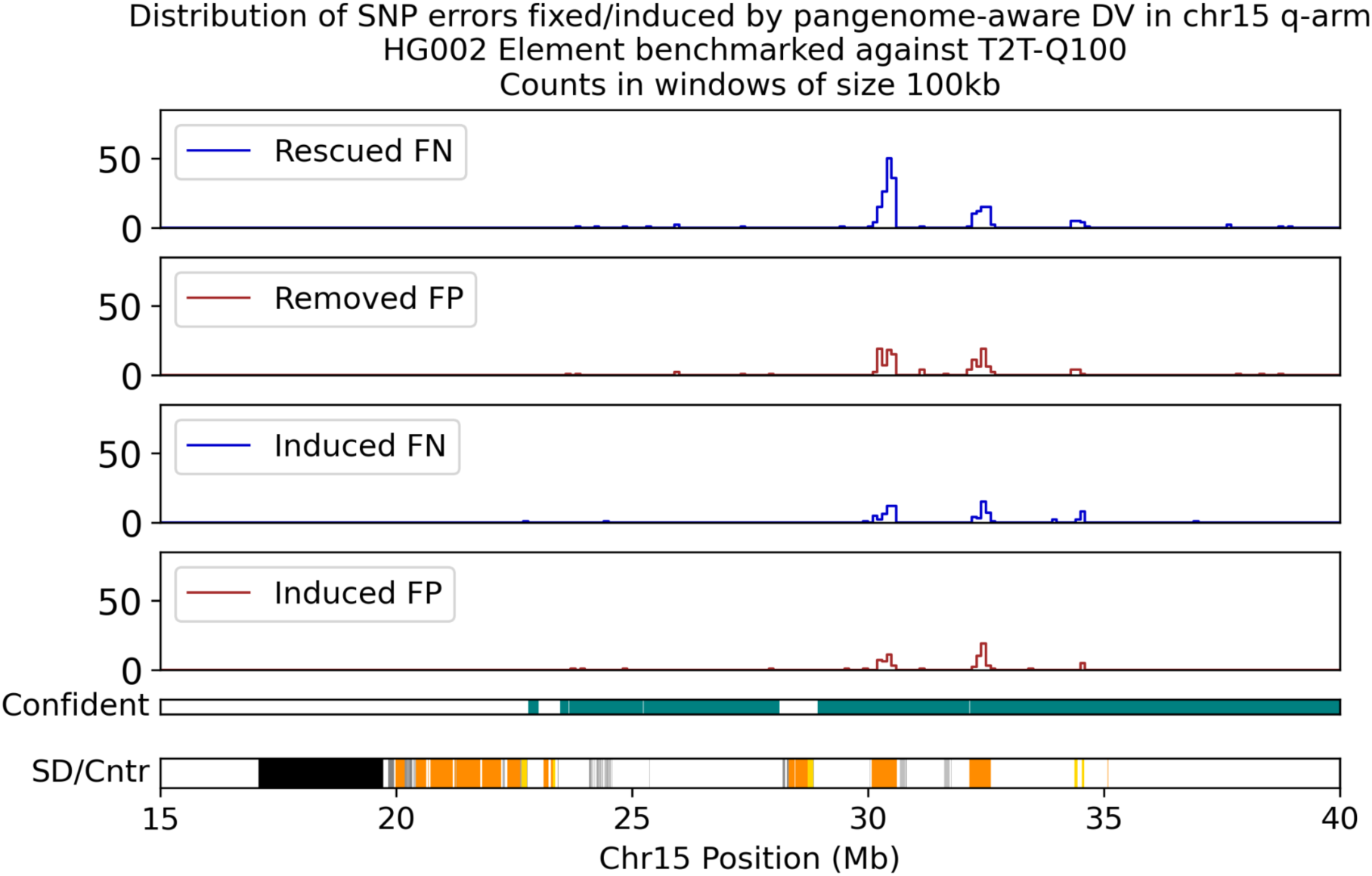
**Locations of SNP errors fixed or induced by pangenome-aware DV in the q-arm of chr15** This figure shows how segmental duplications are enriched with fixed variants (especially for rescued FN in this region). Density of fixed and induced errors are shown in the q-arm of chr15. The number of variants are counted in adjacent non-overlapping windows of length 100kb. Centromere is shown with black color in the bottom track.

**Supplementary Figure 15:**
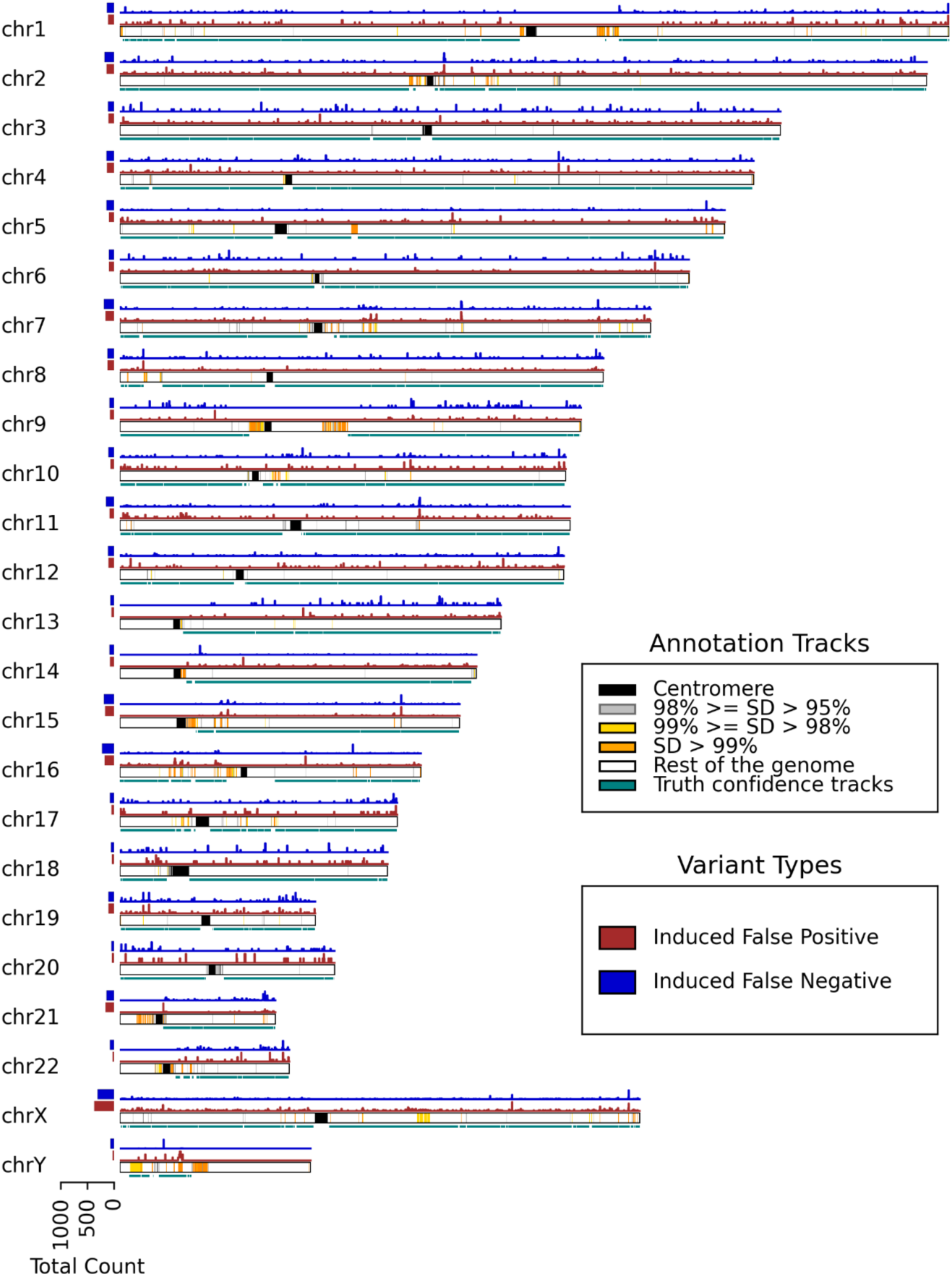
**Ideogram of SNP errors induced by pangenome-aware DV** The locations of the induced errors are plotted across the GRCh38 chromosomes. FP and FN calls are shown with red and blue respectively. The bottom track for each chromosome shows 5 different annotations; centromere (black), SDs with identity greater than 99% (orange), SDs with identity between 99% and 98% (yellow), SDs with identity lower than 95% (light gray) and the rest of the genome (white). Below the SD/Cntr annotation the tracks with teal color show the confidence regions for T2T-Q100 truth set. The left horizontal barplots show the total counts of induced errors per chromosome.

**Supplementary Figure 16:**
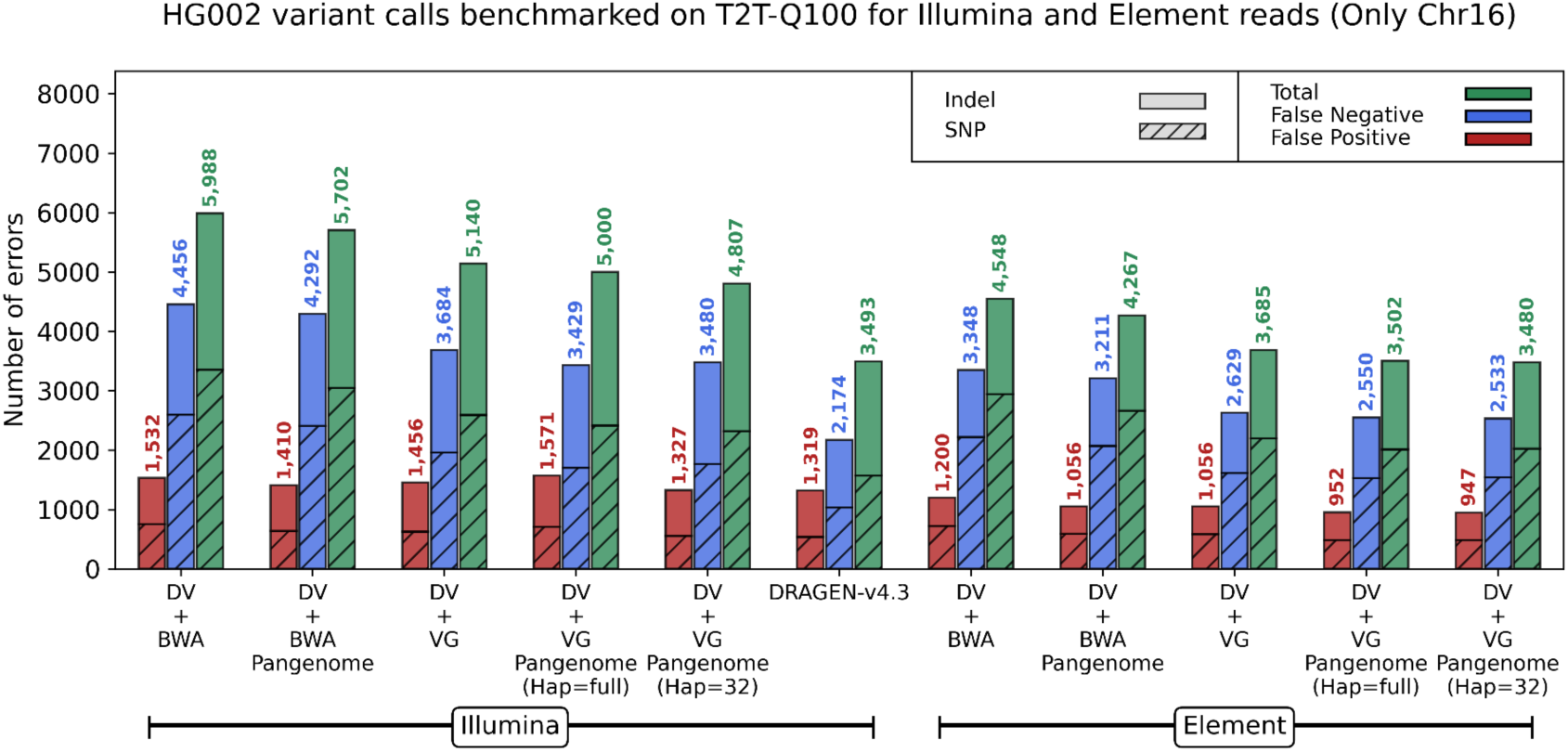
**Benchmarking HG002 variant calls against T2T-Q100 on chr16** HG002 Illumina and Element calls are benchmarked against the T2T-Q100 truth set only on chromosome 16 which was the held-out chromosome for training DeepVariant. Both linear-reference-based DV and pangenome-aware DV have been tested with vg giraffe and BWA-MEM mappers. The x-axis labels with “Pangenome” refer to the pangenome-aware DV and for the rest, the linear-reference-based DV was used for variant calling. The x-axis labels with “(Hap=full)” refers to using all 88 haplotypes in the HPRC- v1.1 pangenome and “(Hap=32)” refers to using a personalized pangenome for HG002 with 32 haplotypes.

**Supplementary Figure 17:**
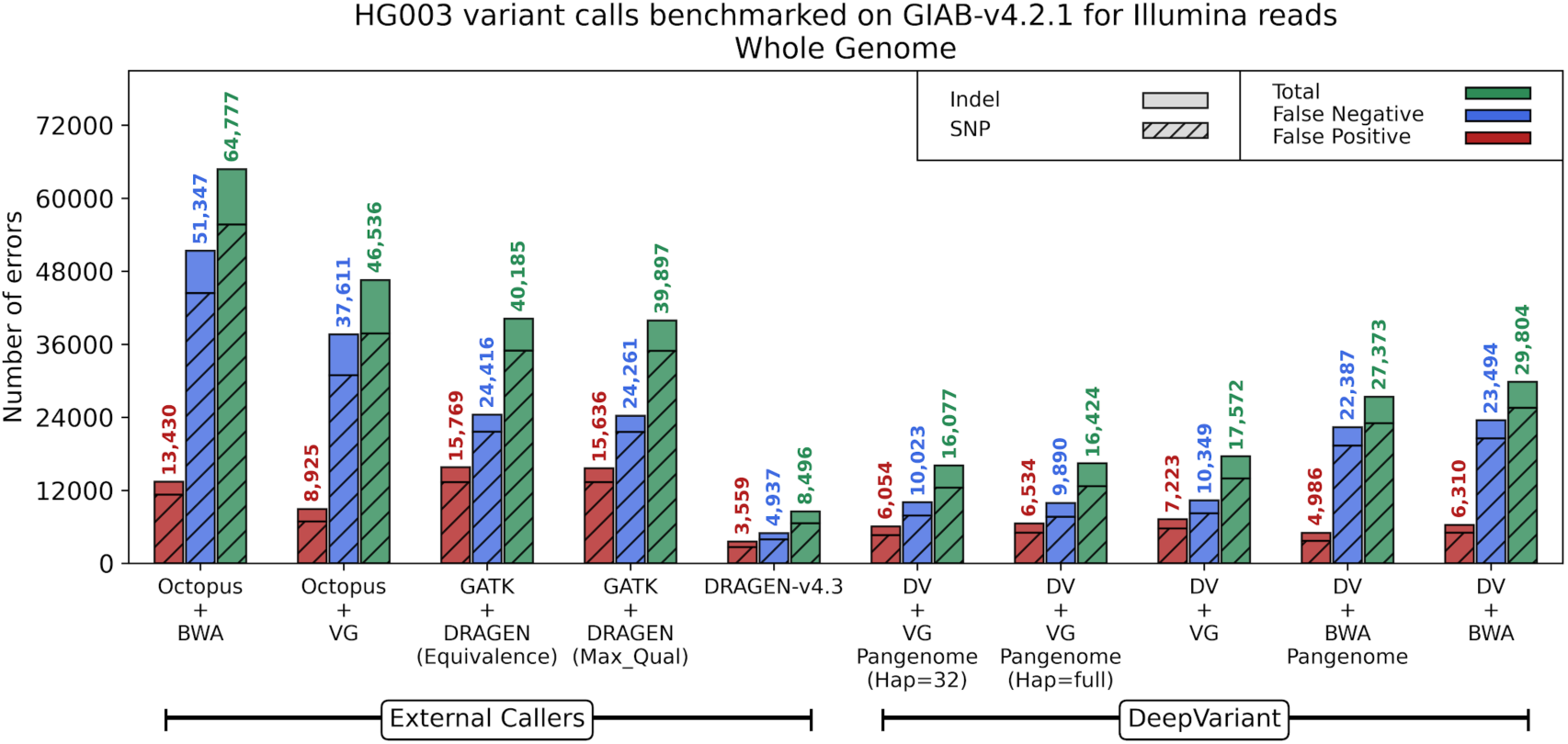
Benchmarking HG003 Illumina calls against GIAB-v4.2.1 truth set. HG003 Illumina calls are benchmarked against the GIAB-v4.2.1 truth set across the whole-genome GIAB high-confidence regions. Different modes of DeepVariant (DV) have been tested with both vg giraffe and BWA-MEM read mappers. The x-axis labels with “Pangenome” refer to the pangenome-aware DeepVariant and for the rest the linear-reference-based DeepVariant was used for variant calling. The x-axis labels with “(Hap=full)” refers to using all 88 haplotypes in the HPRC-v1.1 pangenome and “(Hap=32)” refers to using a personalized pangenome with 32 haplotypes.

**Supplementary Figure 18:**
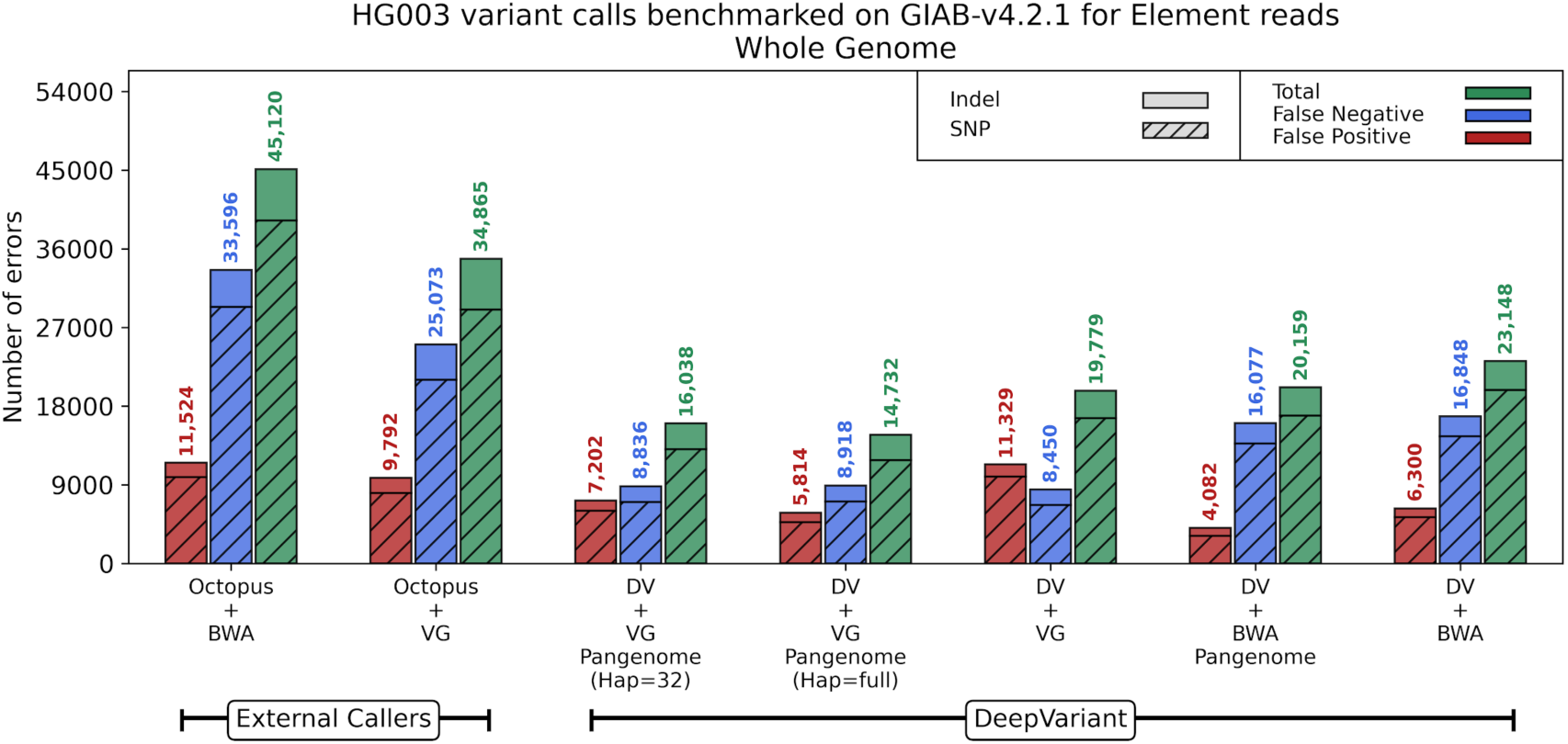
Benchmarking HG003 Element calls against GIAB-sv4.2.1 truth set. HG003 Element calls are benchmarked on the GIAB truth set across the whole-genome GIAB high-confidence regions. Different modes of DeepVariant (DV) have been tested with both vg giraffe and BWA-MEM read mappers. The x-axis labels with “Pangenome” refer to the pangenome-aware DeepVariant and for the rest the linear-reference-based DeepVariant was used for variant calling. The x-axis labels with “(Hap=full)” refers to using all 88 haplotypes in the HPRC-v1.1 pangenome and “(Hap=32)” refers to using a personalized pangenome with 32 haplotypes. The DRAGEN-assisted GATK failed with Element data and Dragen-v4.3 is not designed for Element so they are not reported among the external callers.

## Citations

1. Sherry, S. T. et al. dbSNP: the NCBI database of genetic variation. Nucleic Acids Res 29, 308–311 (2001).

2. Church, D. M. et al. Modernizing Reference Genome Assemblies. PLOS Biology 9, e1001091 (2011).

3. Schneider, V. A. et al. Evaluation of GRCh38 and de novo haploid genome assemblies demonstrates the enduring quality of the reference assembly. Genome Res. 27, 849–864 (2017).

4. Nurk, S. et al. The complete sequence of a human genome. Science 376, 44–53 (2022).

5. Ballouz, S., Dobin, A. & Gillis, J. A. Is it time to change the reference genome? Genome Biology 20, 159 (2019).

6. Magi, A. et al. Characterization and identification of hidden rare variants in the human genome. BMC Genomics 16, 340 (2015).

7. Ebler, J., Schönhuth, A. & Marschall, T. Genotyping inversions and tandem duplications. Bioinformatics 33, 4015–4023 (2017).

8. Pangenomics enables genotyping of known structural variants in 5202 diverse genomes. https://www.science.org/doi/10.1126/science.abg8871 doi:10.1126/science.abg8871.

9. Martiniano, R., Garrison, E., Jones, E. R., Manica, A. & Durbin, R. Removing reference bias and improving indel calling in ancient DNA data analysis by mapping to a sequence variation graph. Genome Biology 21, 250 (2020).

10. Gong, Y., Li, Y., Liu, X., Ma, Y. & Jiang, L. A review of the pangenome: how it affects our understanding of genomic variation, selection and breeding in domestic animals? Journal of Animal Science and Biotechnology 14, 73 (2023).

11. Eizenga, J. M. et al. Pangenome Graphs. Annu Rev Genomics Hum Genet 21, 139–162 (2020).

12. Liao, W.-W. et al. A draft human pangenome reference. Nature 617, 312–324 (2023).

13. Poplin, R. et al. A universal SNP and small-indel variant caller using deep neural networks. Nat Biotechnol 36, 983–987 (2018).

14. Behera, S. et al. Comprehensive genome analysis and variant detection at scale using DRAGEN. Nat Biotechnol 1–15 (2024) doi:10.1038/s41587-024-02382-1.

15. Genomics in the Cloud[Book]. https://www.oreilly.com/library/view/genomics-in-the/9781491975183/.

16. CalculateGenotypePosteriors. *GATK* https://gatk.broadinstitute.org/hc/en-us/articles/360037226592-CalculateGenotypePosteriors (2023).

17. Chen, N.-C. et al. Improving variant calling using population data and deep learning. BMC Bioinformatics 24, 197 (2023).

18. Sibbesen, J. A., Maretty, L., Danish Pan-Genome Consortium & Krogh, A. Accurate genotyping across variant classes and lengths using variant graphs. Nat Genet 50, 1054– 1059 (2018).

19. Ebler, J. et al. Pangenome-based genome inference allows efficient and accurate genotyping across a wide spectrum of variant classes. Nat Genet 54, 518–525 (2022).

20. Li, H. Aligning sequence reads, clone sequences and assembly contigs with BWA-MEM. Preprint at 10.48550/arXiv.1303.3997 (2013).

21. Li, H. & Durbin, R. Fast and accurate short read alignment with Burrows-Wheeler transform. Bioinformatics 25, 1754–1760 (2009).

22. Sirén, J. et al. Personalized pangenome references. Nat Methods 21, 2017–2023 (2024).

23. Zook, J. M. et al. An open resource for accurately benchmarking small variant and reference calls. Nat Biotechnol 37, 561–566 (2019).

24. marbl/HG002. MarBL (2025).

25. Genome in a Bottle. NIST (2012).

26. Kronenberg, Z. et al. The Platinum Pedigree: A long-read benchmark for genetic variants. 2024.10.02.616333 Preprint at 10.1101/2024.10.02.616333 (2024).

27. Carroll, A. et al. Accurate human genome analysis with Element Avidity sequencing. 2023.08.11.553043 Preprint at 10.1101/2023.08.11.553043 (2023).

28. Wang, T. et al. The Human Pangenome Project: a global resource to map genomic diversity. Nature 604, 437–446 (2022).

29. Allendorf, F. W., Hössjer, O. & Ryman, N. What does effective population size tell us about loss of allelic variation? Evol Appl 17, e13733 (2024).

30. Johnston, S. E. Understanding the Genetic Basis of Variation in Meiotic Recombination: Past, Present, and Future. Mol Biol Evol 41, msae112 (2024).

31. Stapley, J., Feulner, P. G. D., Johnston, S. E., Santure, A. W. & Smadja, C. M. Recombination: the good, the bad and the variable. Philos Trans R Soc Lond B Biol Sci 372, 20170279 (2017).

32. A draft human pangenome reference | Nature. https://www.nature.com/articles/s41586-023-05896-x.

33. GBZ file format for pangenome graphs | Bioinformatics | Oxford Academic. https://academic.oup.com/bioinformatics/article/38/22/5012/6731924.

34. Siren, J. jltsiren/gbwtgraph. (2025).

35. Chapter 18. Boost.Interprocess - 1.77.0. https://www.boost.org/doc/libs/1_77_0/doc/html/interprocess.html.

36. Park, J. et al. DeepSomatic: Accurate somatic small variant discovery for multiple sequencing technologies. 2024.08.16.608331 Preprint at 10.1101/2024.08.16.608331 (2024).

37. Kronenberg, Z. et al. The Platinum Pedigree: A long-read benchmark for genetic variants. 2024.10.02.616333 Preprint at 10.1101/2024.10.02.616333 (2024).

38. marbl/HG002. MarBL (2025).

39. McKenna, A. et al. The Genome Analysis Toolkit: A MapReduce framework for analyzing next-generation DNA sequencing data. Genome Res. 20, 1297–1303 (2010).

40. DePristo, M. A. et al. A framework for variation discovery and genotyping using next-generation DNA sequencing data. Nat Genet 43, 491–498 (2011).

41. Whole Genome Germline Single Sample Overview | WARP. https://broadinstitute.github.io/warp/docs/Pipelines/Whole_Genome_Germline_Single_Sample_Pipeline/README/(2025).

42. DRAGEN-GATK Update: Let’s get more specific. *GATK* https://gatk.broadinstitute.org/hc/en-us/articles/360039984151-DRAGEN-GATK-Update-Let-s-get-more-specific (2023).

